# Glucocorticoid signalling exerts distinct lineage-permissive and dose-dependent disruptive actions during human osteoblast differentiation

**DOI:** 10.64898/2026.09.18.752640

**Authors:** Martin Rønn Madsen, Dalia Ali, Mathilde Palmier, Atenisa Caci, Vyacheslav Akimov, Anders Kristian Haakonsson, Blagoy Blagoev, Moustapha Kassem, Susanne Mandrup, Alexander Rauch

## Abstract

At physiological levels glucocorticoid (GC) signalling within the osteoblast lineage has been proposed to be required for skeletal homeostasis, whereas supraphysiological activation supresses bone formation and causes bone loss. While these observations suggest that GC exert dose- and context-dependent effects, the molecular mechanisms underlying these opposing actions remain unresolved. Here we systematically dissected the receptor-, lineage-, and concentration-dependent effects of GC signalling on human bone marrow stromal cell (BMSC) differentiation using integrated transcriptional, epigenomic, and functional analyses. Our findings suggest that GC opposing effects are not simply due to a GR dose-response but instead reflect distinct context- and concentration-dependent modes of action. During early BMSC differentiation, the cellular context induced GR-dependent licensing of lineage-specific gene and enhancer programs. The adverse effects of high-dose GC developed over time and could not be predicted from the early transcriptional response. We found that high-dose GC caused osteoblast maturation arrest, accompanied with a cellular stress response sustained by myeloid zinc finger 1 (MZF1) also identified as an endogenous inhibitor of osteoblast differentiation.

## Introduction

Glucocorticoids (GC) are steroid hormones produced by the adrenal cortex and serve as key endocrine signalling molecules. They regulate a wide range of physiological processes, including skeletal growth and maintenance (Zhou et al., 2013). Due to their anti-inflammatory properties, synthetic analogous have been developed to treat inflammatory and autoimmune diseases, which at high-dose over a prolonged period are associated with severe skeletal complications including glucocorticoid-induced osteoporosis (GIO) and fragility fractures (Hofbauer et al., 2025).

Genetic and pharmacological studies in mice showed that GC signalling within the osteoblast lineage is required for skeletal homeostasis at physiological levels (Macfarlane et al., 2026; Pierce et al., 2019; Rauch et al., 2010; Sher et al., 2006; Sher et al., 2004), whereas supraphysiological activation supresses bone formation and causes bone loss (O’Brien et al., 2004; Rauch et al., 2010). The detrimental effects of GC excess are well supported by clinical observations in patients receiving pharmacological GCs (van Staa et al., 2000) and in patients with endogenous hypercortisolism (Zdrojowy-Welna et al., 2026). In contrast, human conditions associated with impaired glucocortcioid signalling provide limited insight into its physiological requirement in bone. GC resistance caused by mutations in *NR3C1* (Kino et al., 2000), or impaired cortisol regeneration resulting from mutations in *HSD11B1* and *H6PD* (Draper et al., 2003; Lavery et al., 2008; Lawson et al., 2011), are accompanied by compensatory endocrine adaptations and lack a consistent skeletal phenotype. Nevertheless, profound osteoporosis has been reported in a patient carrying a frameshift mutation (p.Ile465Serfs*22) in *NR3C1* (Al Argan et al., 2018) and patients with Addison’s disease have an increased risk of osteoporosis and fractures independent of GC replacement dose (Stergianos et al., 2025). Thus, while human studies firmly establish the detrimental skeletal effects of GC excess, a requirement for physiological GC signalling in human bone remains unresolved. Interestingly, *in vitro* model systems studying the differentiation of human osteoblasts traditionally require the addition of dexamethasone (Della Bella et al., 2021; Langenbach & Handschel, 2013; Mollentze et al., 2021), a synthetic GC analog that is likewise used as part of adipogenic induction cocktails to drive fat cell differentiation (Scott et al., 2011). Thus, GC can exert both permissive and inhibitory actions on osteoblast differentiation, supporting osteogenic differentiation under some conditions while inhibiting bone formation under others, suggesting that their effects on osteoblasts are determined not only by dose but also by signaling context.

The transcriptional response to glucocortcioids is strongly dependent on cell-type identity. Hormone-induced GR binding is largely constrained by chromatin accessibility (John et al., 2011), with cell-type specific GR binding preferentially occuring at distal enhancers with corresponding cell-type specific accessibility (Love et al., 2017). Within accessible chromatin, GR recruitment is further determined by glucorticoid response element (GRE) strength and interactions with lineage- and context-specific transcription factors (Biddie et al., 2011; Helminen et al., 2024). Accordingly, HNF4A facilitates liver-specific GR binding at weak GREs (Hunter et al., 2022), while NF-kB activation redistributes GR occupancy upon inflammatory stimulation (Mostafa et al., 2025). Importantly, GR cooperates with C/EBPβ and other early adipogenic transcription factors to activate enhancers during early chromatin remodeling (Park & Ge, 2017; Siersbaek et al., 2011; Steger et al., 2010). Thus, rather than acting through a fixed set of genomic targets, GR engages regulatory programs determined by chromatin state and transcription factor interactions. Whether GR similarly engages the osteogenic regulatory landscape to facilitate osteoblast differentiation, and how such lineage-specific actions intersect with the concentration-dependent effects of GC siganling, remains poorly understood.

Here we systematically dissected the receptor-, lineage-, and concentration-dependent effects of GC signalling on human BMSC differentiation using integrated transcriptional, epigenomic, and functional analyses. We uncover a dual role of GC in which GR-dependent licensing of lineage-specific gene and enhancer programs enable stromal cell differentiation, while excessive GC exposure engages myeloid zinc finger 1 (MZF1) in a cumulative cellular stress response, leading to osteoblast maturation arrest.

## Material & Methods

### Reagents

Antibodies, oligo sequences and catalogue numbers of all reagents are outlined in Supplementary table 1.

### Cell culture and differentiation

Telomerase-immortalized human mesenchymal stromal cells (MSC) isolated from the bone marrow (BM-hMSC-TERT4) were generated as previously described (Simonsen et al., 2002). The BM-hMSC-TERT4 cells were used at passages 43 to 50 and grown under standard cell culture conditions (37°C with 5% CO_2_ and 95% humidity) in α-MEM supplemented with 10 % fetal bovine serum (FBS) and 1 % penicillin/streptomycin (P/S). hBMS-TERT4 cells were expanded at a split-ratio of 1:4 twice a week or set out for experiments by trypsinization using 0.05% trypsin–EDTA when reaching 80 to 90% confluency. Contamination with *Mycoplasma* was checked in the research unit every second month with the N-GARDE Mycoplasma PCR Reagent set (Euroclone) using either fresh conditioned media or media stored at 4 °C. All cultures reported here were negative of *Mycoplasma*.

For differentiation, two days post confluency (day 0) cells were induced to undergo osteoblast or adipocyte differentiation by exposing them to differentiation cocktails. For osteoblast differentiation cells were switched to α-MEM supplemented with 10 % FBS, 1 % P/S, 10 nM dexamethasone, 10 nM vitamin D, 10 mM β-glycerophosphate, and 50 µg/ml ascorbic acid. For adipocyte differentiation, cells were switched to DMEM supplemented with 10 % FBS, 1 % P/S, 100 nM dexamethasone, 10 µg/mL insulin, 500 µM IBMX, and 1 µM rosiglitazone. Media was replaced on day 2, 4, 7, 9 and 11.

### Osteogenic and adipogenic functional assays

For Alizarin Red and Oil Red O, cells were washed with PBS and fixed at the indicated time points with 70% ice-cold ethanol for 1hour at -20 °C or 4% formaldehyde in PBS for 30 minutes, respectively. Staining was done with either Alizarin Red (1% solution in water; pH=4.2) or Oil Red O (300mg/mL Oil Red O in isopropanol diluted 60:40 v/v in water) for 30 minutes followed by washing steps with water. Dishes were either scanned or wells were photographed using a 10x phase contrast objective. For quantification, Alizarin Red dye was extracted with water containing 20% methanol and 10% acetic acid, while Oil Red O dye was extracted with isopropanol following measurement of absorbance at 570 and 520 nm on a FLUOstar Omega plate reader, respectively.

To visualize alkaline phosphatase activity, we used the Alkaline Phosphatase Detection Kit (Sigma-Aldrich) on cells which were washed once with PBS and then fixed in acetone/citrate solution (3:2). Cells were incubated with ALP substrate solution (mixture of Fast Violet B Salt and naphthol AS-MX phosphate) for 1 hour at room temperature. Wells were washed twice with PBS and scanned or photographed using a microscope with 10x phase contrast objective. For quantitative measurements, cells were fixed using 4% paraformaldehyde in 90% ethanol for 30 seconds at room temperature; fixative was removed and cells were incubated with p-nitrophenyl phosphate solution (1 tablet in 5 ml water) for 20–30 minutes in the dark at room temperature. Reaction was subsequently stopped by adding 1/3 volume of 3M NaOH, and the ALP activity was reported as absorbance at 405 nm on a FLUOstar Omega plate reader normalized to cell viability.

### Cell viability assay

Cell viability was measured using CellTiter-Blue assay (Promega) by incubating the cells with a mixture of cell viability solution and media (1:5) at 37 °C for 1 h prior to any fixation. Fluorescence intensity (Ex 579 / Em 584) was measured with a FLUOstar Omega plate reader.

### siRNA and shRNA knockdown

For transient transfection, reverse transfection was applied using Lipofectamin™ 2000 (ThermoFischer) and siRNAs Silencer™ Select (Thermo Fischer) at a final concentration of 20 nM. Cells were plated at a density of approximately 50,000 cells per cm^2^ and treated with transfection mix (20 nM siRNA and 0.1% Lipofectamin™ 2000 in Opti-MEM™) for 6 h. Cells reached confluency the day after transfection (day 2 of differentiation). Knockdown efficiency was tested with western blot on day 0 of differentiation.

For stable *MZF1* knock-down, MZF1-specific short hairpin RNA (shRNA) was generated by annealing complementary DNA oligonucleotides containing the shRNA target sequences and hairpin linkers acquired from Integrated DNA Technologies (IDT, Leuven, Belgium). A previously published scrambled shRNA sequence with no target specificity was used as control (Wiebe & Traktman, 2007). The annealed shRNA oligonucleotides, together with an *EEF1A1* promoter-driven puromycin resistance cassette (from pCDF1-MCS2-EF1-Puro-System Biosciences Co.) were cloned into the lentiviral vector pSicoR (Addgene Plasmid 11579) (Ventura et al., 2004), and constructs were verified by DNA sequencing. Generation of lentiviral particles, infection, selection, and monitoring of stable BM-hMSC-TERT4 cells expressing the respective shRNA constructs were performed as previously described (Akimov et al., 2014). Protein expression level of *MZF1* in mutant cells was tested by western blot analysis.

### Western blot

Cultured cells were washed in cold PBS, and protein lysates were prepared using M2 lysis buffer (20 mM Tris pH 7.0; 0.5% NP-40; 250 mM NaCl; 3mM EDTA; 3mM EGTA) including cOmplete™ protease inhibitors (Roche). Lysates were centrifuged at 12,000 g at 4 °C for 10 minutes and protein concentration was determined using Pierce™ BCA Protein Assay Kit (ThermoFischer) and 25-30 µg protein were mixed with 4x NuPAGE™ LDS Sample Buffer and 10x NuPAGE™ Sample Reducing Agent (ThermoFischer) before running on a 10 % NuPAGE™ Bis-Tris Mini Protein Gel (Invitrogen). Blotted nitrocellulose membranes were blocked (5 % BSA in PBS with 0.1 % TWEEN^®^ 20) for 1 hour and subsequently incubated with primary antibodies (1:1,000 in blocking buffer) overnight at 4 °C. Membranes were washed 3 times with PBS 0.1% TWEEN^®^ 20 and incubated with secondary antibodies (1:2,000 in blocking buffer) for 1h at room temperature. After washing three times with PBS 0.1 % TWEEN^®^ 20, membranes were developed by incubating with Pierce™ ECL Western detection substrate (ThermoFischer) for 1 to 5 minutes and immunocomplexes were detected by enhanced chemiluminescence on a ChemiDoc machine.

### Senescence-associated β-galactosidase activity

To examine the cellular senescence, we used the Senescence β-galactosidase Staining and Activity Assay Kit (Cell Signalling) as previously described (Dzubanova et al., 2025). For staining, cells were washed with PBS, fixed for 10 minutes at room temperature and rinsed again with PBS before incubation with the β-galactosidase staining solution (pH = 6.0) at 37 °C in dry incubator (no CO_2_) overnight. 6-12 hours post-incubation, images of blue color senescent cells were taken with a 10x phase contrast objective. For quantitative measurement, lysates were treated according to manufacturer’s instructions and fluorescence intensity (Ex 360 / Em 465) was measured with a FLUOstar Omega plate reader.

### Bioenergetic assay

Cell bioenergetic analysis was performed using Seahorse XFe96 extracellular flux analyzer (Agilent) to measure the Oxygen Consumption Rate (OCR) and Extracellular Acidification Rate (ECAR) simultaneously. Cells were plated at a density of 1,000 cells per well in a Seahorse XFe96 cell culture microplate (96 wells, Agilent) in 120 µL of growth media. The cells were either left undifferentiated for 12 days, or osteogenic differentiation with 10 or 100 nM Dex was initiated sequentially 12, 7 and 1 day before analysis. The medium was changed every 2 to 3 days.

On the day of the experiment, the assay medium was prepared by supplementing Seahorse XF DMEM medium (Agilent) with 10 mM glucose, 2 mM Glutamine, and 1 mM Sodium Pyruvate. An hour before the measurement, the cell culture medium was replaced with the assay medium and incubated in a non-CO_2_ environment at 37°C. For the analysis, we used the Seahorse XF T Cell Metabolic Profiling Kit (Agilent) following the manufacturer instructions. In brief, the first injection was ATP-synthetase inhibitor Oligomycin (1.5 µM final/well), the second injection was mitochondrial membrane uncoupler BAM 15 (2.5 µM final/well), the third injection was a mix of complex I and III inhibitors Rotenone/Antimycin A (0.5 µM each final/well). Four to six wells per condition were used.

After OCR and ECAR measurements were completed, we added Hoechst 33342 (Invitrogen) to each well as nuclear staining (20 µM / final well) for 15 minutes in the dark at 37°C. The cells were imaged using a Cytation1 (Biotek, Agilent) and the number of nuclei in each well was counted using Cell Imaging software (Agilent). The OCR and ECAR values were normalized to 1,000 cells in Wave Software (Agilent).

### Ectopic implantation

To assess bone-forming capacity we used ectopic transplantation in 14-week-old immune deficient female mice as previously described in detail (Abdallah et al., 2008). We made 4 subcutaneous cavities, of which 3 were used for implants and 1 for prednisolone or placebo slow-release pellets, resulting in a calculated dose of 6.25 mg/kg/day (7.5 mg; 60-day release; Innovative Research of America, Inc.). We used 500,000 cells per implant which were harvested after 8 weeks and fixed in 4 % PFA for 24 h followed by decalcification in 0.5 mol/L formic acid for 72 h. Implants were paraffine-embedded and sectioned at three different depths (100 µm apart). H&E-stained sections were quantified for bone using ImageJ by marking areas of bone normalized to the total area of the implant. Average of the three depths was used to report bone area / tissue area. As a deviation from the published protocol, NOD.Cg-*Prkdc^scid^ Il2rg^tm1Sug^*/JicTac (Taconic) mice have been used instead of NOD/LtSz-*Prkdc*^scid^ mice. Ethical approval was given by the Animal Experimentation Council of the Ministry of Food, Agriculture and Fishing Denmark (License 2022-15-0201-01225).

### RNA extraction

Cells were rinsed with ice-cold PBS and harvested in Trizol (Sigma-Aldrich). After adding 0.2 volume chloroform, tubes were vortexed for 30 seconds, rested for 15 minutes at room temperature, and were centrifuged at 10,000g, at 4 °C for 10 minutes. The upper phase was transferred to a new tube, and 0.6 volume of ethanol was added. After mixing, the sample was loaded on a EconoSpin^®^ column and washed 3 times with 500 µL RPE buffer (Qiagen) with spinning at 13,000 g for 1 minute at room temperature. RNA was eluted with 30 µL of DEPEC-treated water. RNA concentration and quality was determined with a Nanodrop and the RNA Kit (Agilent) on a Fragment Analyzer.

### cDNA synthesis and RT-qPCR

For RT-qPCR analysis, 1 µg RNA was denaturized and reversely transcribed using the RevertAid First Strand cDNA Synthesis Kit (ThermoFischer). Quantitative RT-qPCR was performed on Biorad CFX connect RT-qPCR machine using Fast SYBR™ Green Master Mix (Applied Biosystems). *ACTB* (β-actin) was used to normalize gene expression.

### RNA-sequencing

Unstranded RNA-seq libraries were constructed according to manufacturer’s instructions (TruSeq 2, Illumina) using 1 µg of total RNA. Libraries were sequenced on an Illumina Novaseq and reads were mapped to the human genome (GRCh37) using Spliced Transcripts Alignment to a Reference (STAR) (Dobin et al., 2013). Tag counts were summarized at the gene level using HOMER (Heinz et al., 2010) allowing only one read per position per length. Differential gene expression analysis and gene count transformation and normalization were done with DESeq2 (Love et al., 2014). Differentially expressed genes were defined using a threshold of lower than 0.01 for the adjusted p-value (p-adj < 0.01). Gene ontology (Ashburner et al., 2000) and Reactome pathway (Ragueneau et al., 2026) enrichment analysis was performed using goseq (Young et al., 2010).

To rank genes for fgsea (bioRxiv doi:10.1101/060012) based gene set enrichment analysis we used the DEseq2-derived Wald statistic (lfcSE) leveraging effect and uncertainty in gene regulation.

### Enhancer mapping

DNase-seq, MED1 ChIP-seq and enhancer coordinates were used from our previously published enhancer activity mapping through osteoblast and adipocyte differentiation of hBM-MSC-TERT4 cells (Rauch et al., 2019). H3K27ac ChIP-seq was performed as previously described (Nielsen & Mandrup, 2014; Siersbaek et al., 2014). Briefly, BM-hMSC-TERT4 cells were harvested on day 0, and day 1 after induction with osteogenic or adipogenic components and the indicated concentrations of dexamethasone. Cells were cross-linked on the culture dishes for 10 minutes in 1 % formaldehyde in PBS followed by quenching with 0.125 M glycine. Cells were scraped off, spun down at 13,000g and following shearing of chromatin on a Bioruptor (Diagenode), immunoprecipitation was performed on material of approximately 0.5 million cells using H3K27ac antibody (Abcam). Approximately 10-20 ng of immunoprecipitated DNA was prepared for sequencing according to the instructions by the manufacturer (Illumina).

Sequencing reads were mapped to the human genome (GRCh37) using STAR (Dobin et al., 2013). BEDTools (Quinlan & Hall, 2010) was used to expand previously defined enhancer regions (Rauch et al., 2019) by 150 % to quantify H3K27ac tag count density and to link enhancer regions with transcription start sites within a window of 50 kb. Sequencing tags allowing only one read per position per length and GRE motif score in peaks were quantified using HOMER (Heinz et al., 2010). Differential occupancy and tag density normalization was done using DEseq2 (Love et al., 2014). Dynamic enhancers were defined using a threshold of lower than 0.05 for the adjusted p-value (p-adj < 0.05).

### Motif activity

Motif activity and target gene prediction were either based on gene expression alone (Fastq files) using the ISMARA tool (Balwierz et al., 2014), or on gene expression and condition matching H3K27ac ChIP-seq data (DESeq2 normalized tag counts and enhancer coordinates) using the IMAGE tool (Madsen et al., 2017). Motifs with significant changes were determined using the ISMARA derived z-score a threshold of greater than 2, while IMAGE based significance was determined using a paired t-test and p-value threshold lower than 0.05. For network construction, both target genes and motifs were restricted to genes present in our gene count matrix. Target genes were compared to all detected genes using Wald stat ranking-based gene set enrichment analysis or a Wilcox test based on absolute fold changes.

### Statistics and reproducibility

All statistical analysis were carried out in R, and unpaired student’s t-test was used for RT-qPCR data and quantification of ALP activity, matrix mineralization, lipid droplet formation, β-galactosidase activity, and seahorse parameters in hBMSC-TERT4 cells. Paired student’s t-test was used for implantation assays to account for implant location effects. Statistics for sequencing-based methods were derived from DEseq2, goseq, fgsea, or a wilcox test. Replicates and statistical tests are outlined in the individual figures. For effect size quantification we used the Cohen’s d (variation around the mean) or Cliff’s delta (for ranked tests) from the effsize package (zenodo doi: 10.5281/zenodo.1480624) in R.

## Results

### hBMSC differentiation is sensitive to GC concentration and mediated through GR

Osteogenic and adipogenic *in vitro* differentiation protocols employ distinct glucocorticoid (GC) concentrations, raising the question of how GC signalling contributes to differentiation outcomes in a lineage- and concentration-dependent manner. Using telomerase-immortalized human bone marrow derived stromal cells (hBMSC-TERT4-cells) (Simonsen et al., 2002), we verified GC signalling to be indispensable for matrix mineralization during osteogenic (Fig. 1A) and lipid accumulation during adipogenic differentiation (Fig. 1B). While differentiation was poor in the absence of dexamethasone (Dex), adipogenic differentiation was enhanced in a dose dependent manner (Fig. 1B), and osteogenic conversion was suppressed at high GC concentration (Fig. 1A). Osteoblast differentiation was optimal at 10 nM Dex while adipocyte formation was best at 100 nM Dex. The lineage and concentration dependent effects of GC on MSC differentiation can be recapitulated by substituting Dex with the natural GC cortisol but not aldosterone which activates the mineralocorticoid receptor (Fig. 1A, B). Pharmacological inhibition of the glucocorticoid receptor (GR) using mifepristone abolished Dex-induced osteogenic (Fig. 1C) and adipogenic differentiation (Fig. 1D). Consistently, genetic depletion of GR via *NR3C1* knockdown (Fig. 1E & Supplementary Fig. 1A) markedly impaired both mineralization and lipid accumulation (Fig. 1F). In sum, these findings establish GR signalling as an essential requirement for both osteoblast and adipocyte differentiation in human stromal cells.

**Figure 1.**
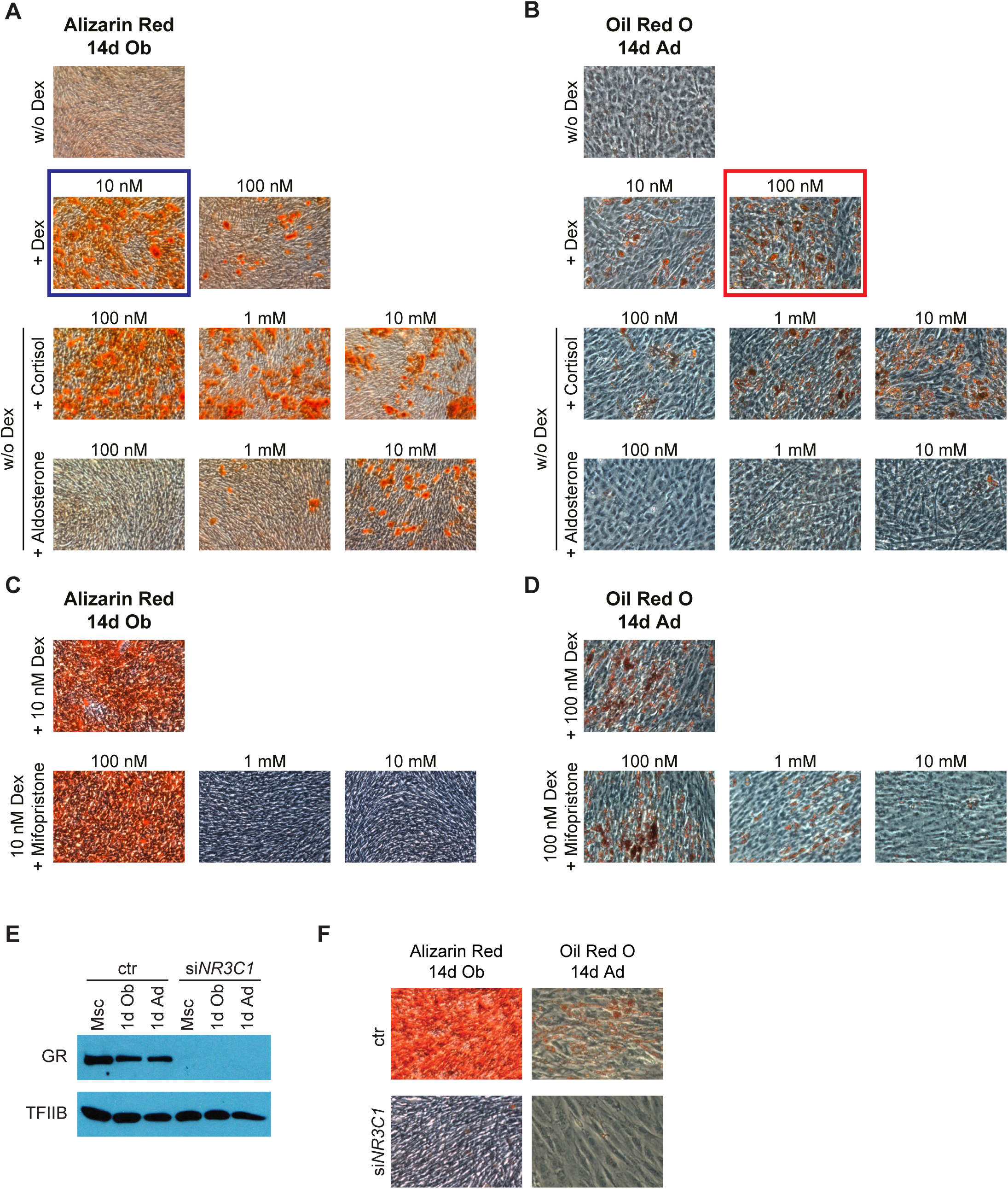
Glucocorticoids stimulate osteoblast and adipocyte differentiation in human stromal cells. (**A**) Alizarin Red staining of hBM-MSC-TERT4 cells after 14 days of osteogenic differentiation with the indicated concentrations of dexamethasone (Dex), cortisol, and aldosterone. The blue box highlights the standard condition for osteoblast differentiation with hBM-MSC-TERT4 cells. (**B**) Oil Red O staining of hBM-MSC-TERT4 cells after 14 days of adipogenic differentiation with the indicated concentrations of Dex, cortisol, and aldosterone. The red box highlights the standard condition for osteoblast differentiation with hBM-MSC-TERT4 cells. (**C**) Alizarin Red staining of hBM-MSC-TERT4 cells after 14 days of osteogenic differentiation with 10 nM Dex and the indicated concentrations of mifopristone. (**D**) Oil Red O staining of hBM-MSC-TERT4 cells after 14 days of adipogenic differentiation with 100 nM Dex and the indicated concentrations of mifopristone. (**E**) Western blot analysis of GR and TF2B in hBM-MSC-TERT4 cells with *NR3C1* knockdown or control siRNA. Protein was harvested 72 hours after siRNA transfection in cells exposed to growth, osteogenic, or adipogenic media for 24 hours. (**F**) Alizarin Red (left) and Oil Red O (right) of hBM-MSC-TERT4 cells after 14 days of osteogenic and adipogenic differentiation. Cells were transfected with control or *NR3C1* targeting siRNA three days prior to induction of differentiation.

### GC signalling is required to unlock lineage-specific gene programs with a strong effect on osteogenic commitment

To investigate how GR signalling shapes early lineage commitment, we performed RNA-seq one day after induction of osteogenic or adipogenic differentiation in hBMSC-TERT4-cells in the presence or absence of GC signalling using different doses of Dex or genetic ablation of GR (*NR3C1* gene). Principal component analysis suggested that Dex withdrawal (Fig. 2A) and GR ablation (Fig. 2B) similarly impair the establishment of early lineage-selective transcriptional states. Osteogenically stimulated samples in the absence of Dex or GR largely overlapped with undifferentiated cells, indicating a particularly strong requirement for GC signalling to establish the early osteoblast gene program. GC and GR signalling were likewise important for the activation of canonical GR target genes (*FKBP5*, *DUSP1*, and *ZBTB16*) as well as genes with osteoblast (*CRYAB*, *SCUBE3*, and *OMD*) and adipocyte-selective (*ACSL1*, *CEBPA*, and *TMEM64*) induction (Fig. 2C). In line, early regulation of genes linked to osteoblast and adipocyte function was abolished in the absence of Dex or GR signalling (Supplementary Fig. 2A).

**Figure 2.**
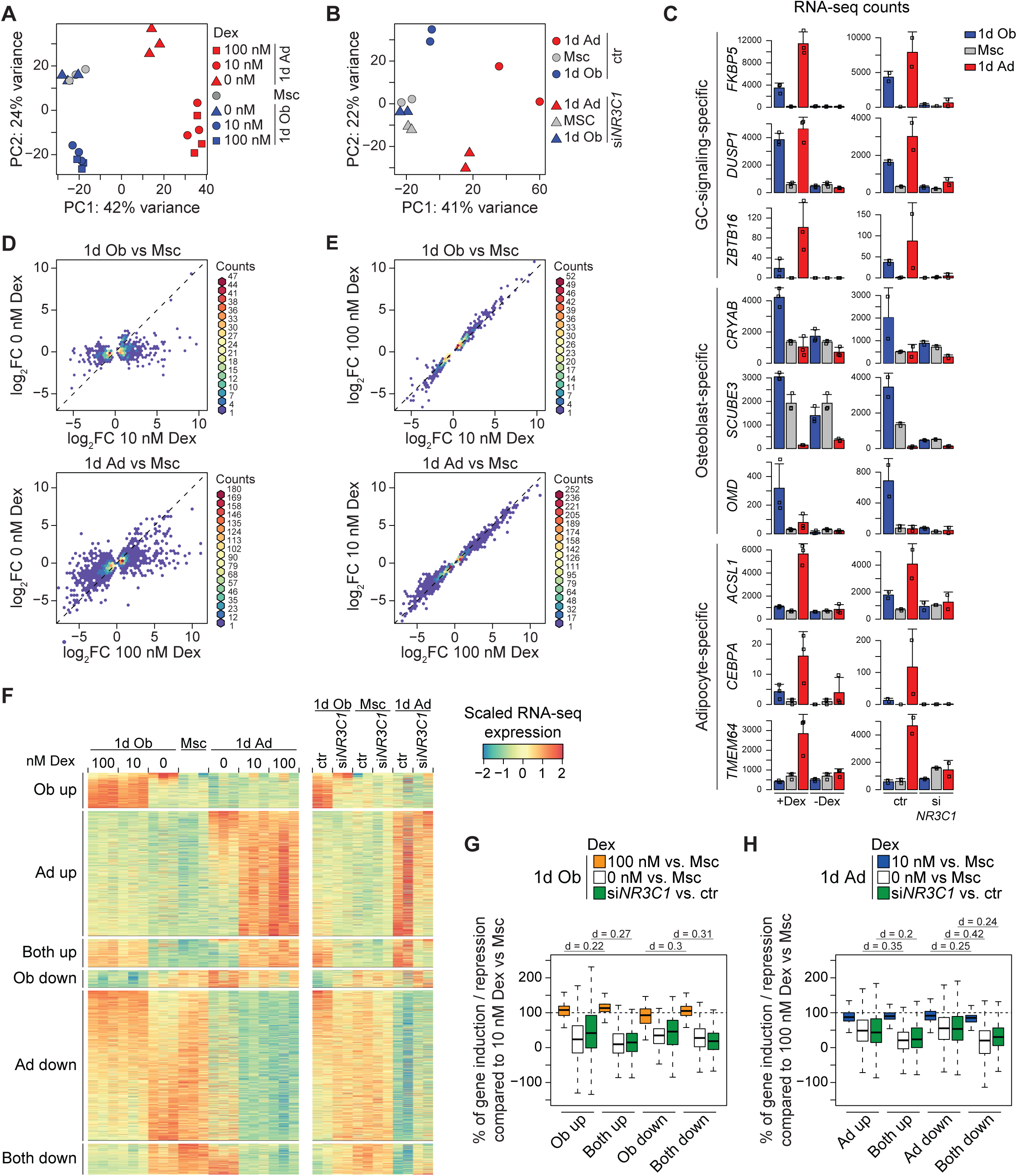
Glucocorticoid action drives early lineage-selective genes in a dose-independent manner. (**A**) PCA plot of RNA-seq data from hBM-MSC-TERT4 cells prior to and after day 1 of differentiation in the presence and absence of Dex. (**B**) PCA plot of RNA-seq data from hBM-MSC-TERT4 cells prior to and after day 1 of differentiation. Cells were treated with control or *NR3C1* targeting siRNA 3 days prior to induction of differentiation. (**C**) Bar plots showing RNA-seq based gene counts of osteoblast-specific, adipocyte-specific, and commonly upregulated genes in hBM-MSC-TERT4 cells on day 1 of differentiation that were affected by loss of GC signalling. Cells were either differentiated in the presence and absence of Dex (left panel, n = 3) or treated with control or *NR3C1* targeting siRNA three days prior to differentiation (right panel, n = 2). Bar showing mean and error bars depicting standard deviation. (**D**) Density scatter plot comparing changes in gene expression during 1 day of osteoblast (upper panel) and 1 day of adipocyte differentiation (lower panel) with and without Dex. Genes selected on differential expression (DESeq2 padj < 0.01) during standard conditions. (**E**) Density scatter plot comparing changes in gene expression during 1 day of osteoblast (upper panel) and 1 day of adipocyte differentiation (lower panel) with 10 and 100 nM of Dex. Genes selected on differential expression (DESeq2 padj < 0.01) during standard conditions. (**F**) Heat map illustrating osteoblast and adipocyte-specific as well as commonly induced or repressed gene expression upon 1 day of osteoblast or adipocyte differentiation in hBM-MSC-TERT4 cells with indicated concentrations of Dex (left panel) or upon control or *NR3C1* knockdown (right panel). Genes are ordered similarly in both heat maps based on significant changes (DESeq2-padj < 0.01) during standard conditions (left). Genes can be assigned to induced and repressed groups simultaneously. (**G**) Box plot (band, median; box, first and third quartiles; whiskers, 1.5 times the interquartile range) quantifying gene regulation on day 1 of osteoblast differentiation, normalized to the respective gene regulation with standard osteoblast differentiation (10 nM Dex or control siRNA; 100%). Gene groups are based on Figure 1F. Effect sizes as Cliff’s delta. (**H**) Box plot (band, median; box, first and third quartiles; whiskers, 1.5 times the interquartile range) quantifying gene regulation on day 1 of adipocyte differentiation, normalized to the respective gene regulation during standard adipocyte differentiation (100 nM Dex or control siRNA; 100%). Gene groups are based on Figure 1F. Effect sizes as Cliff’s delta.

While addition of Dex had a huge impact on early gene regulation (Fig. 2D), there was only a minor amplification of regulation when comparing high (100 nM) versus low (10 nM) Dex concentrations (Fig. 2E) on day 1 of differentiation. In relation to the number of genes regulated during osteoblast and adipocyte differentiation, we found GC signalling to be more instrumental for osteoblast than adipocyte differentiation (Supplementary Fig. 2B & C). To further resolve how GC signalling interacts with lineage context, we analysed early gene regulation grouping gene induction and repression depending on if they were common or specific to a lineage (Fig. 2F), for which we found almost mirroring effects comparing ligand and receptor deprivation. When deciphering lineage- and GC concentration-effects on gene regulation, we found genes regulated in both lineages to be most sensitive to GC concentration and GR ablation during both early osteoblast (Fig. 2G) and adipocyte differentiation (Fig. 2H). Among lineage-selective genes, however, loss of induction or repression following reduction of GC signalling was more pronounced for the osteoblast than the adipocyte program, despite osteoblast differentiation requiring the lowest Dex concentration. This difference could be quantified when comparing the lineage-selective responses at equal GC concentrations (Supplementary Figure 2D & E). Together, these results demonstrate that GC signalling acts in a permissive manner to allow lineage-specific transcriptional programs with GR activity being more important for osteogenic than adipogenic gene regulation. Thus, while GC signalling is required to unlock lineage-specific gene programs, GC concentration primarily tunes transcriptional amplitude rather than determining lineage identity.

### GC signalling controls lineage-selective enhancer activation especially during early osteogenic differentiation

To determine how GR signalling interfaces with chromatin regulation during early stromal cell differentiation, we integrated Dex-dependent profiles of GR binding and enhancer acetylation (H3K27ac) with previously established enhancer marks during early differentiation of hBMSC-TERT4 cells (Rauch et al., 2019).

Genome browser inspection revealed lineage-dependent enhancer activation and GR recruitment to loci associated with osteoblast-specific, adipocyte-specific, and commonly regulated genes (Fig. 3A). At a genome wide level, differentiation in the presence of Dex led to widespread changes in enhancer acetylation (Fig. 3B and Supplementary Fig. 3A & B), which was only slightly increased with hormone dose (Fig. 3C), in line with the observations on gene regulation. To link GC-dependent enhancer activation to transcriptional outcomes, we observed a local enrichment of enhancers and genes with strong GC dependency for both GC-mediated induction and repression (Fig. 3D) as well as an enrichment of eQTL in coherent gene and enhancer clusters (Supplementary Fig. 3C). With focus on GR occupancy, we found GR binding levels to be highly predictive for enhancer activation (Fig. 3E), which was further reflected by GR recruitment to enhancers with osteoblast-specific, adipocyte-specific, and common gain of H3K27-acetylation (Fig. 3F). Importantly, for all activated enhancer groups, H3K27ac-levels (Fig. 3G) and GR binding (Fig. 3H) were strongly induced by GC-signalling with a minor amplification by increasing hormone concentrations. Similar to gene regulation, commonly induced enhancers showed the strongest GC-sensitivity (Fig. 3G and Supplementary Fig. 3D & E), while osteoblast-specific gain in enhancer activity was more dependent on GC-signalling (Fig. 3G) and more evident when normalizing to similar Dex concentrations (Supplementary Fig. 3F & G). GR occupancy increased with rising Dex concentrations across all activated enhancer groups and in both differentiation conditions (Fig. 3H). While commonly activated enhancers showed comparable GR binding intensities for equal Dex concentrations, lineage-selectively activated enhancers preferentially recruited GR in the corresponding differentiation condition. However, the lineage preference and concentration effect equalized at osteogenic induced enhancers, which showed indistinguishable GR binding during standard adipocyte and osteoblast differentiation (Fig. 3H). Thus, GR-binding seems instrumental for enhancer-activation but is not sufficient to explain the lineage-selective enhancer activity landscape during early osteoblast and adipocyte differentiation.

**Figure 3.**
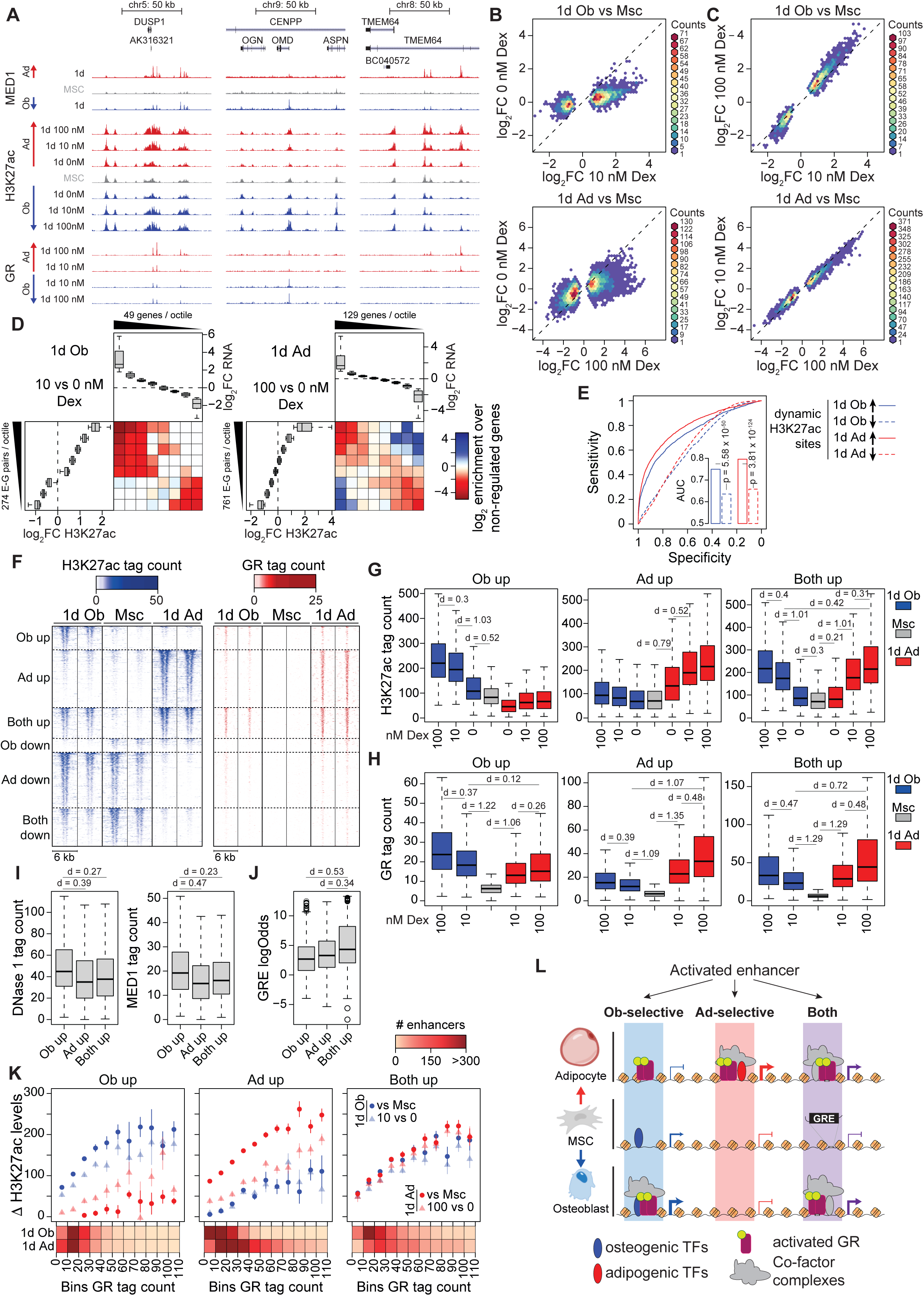
GR acts as instructive chromatin licensing factor in a lineage but not in a dose-dependent manner during early stromal cell differentiation. (**A**) UCSC genome browser screen shot showing ChIP-seq tag density for MED1 and GR recruitment as well as H3K27-acetylation in hBM-MSC-TERT4 cells prior to and after one day of osteoblast and adipocyte differentiation. Dex concentrations are depicted. Loci of *DUSP1*, *OMD*, and *TMEM64* as examples of commonly, osteoblast, and adipocyte-specific induced genes. (**B**) Density scatter plot of H3K27ac ChIP-seq tag counts in hBM-MSC-TERT4 cells on day 1 of osteoblast (upper panel) or adipocyte differentiation (lower panel) with and without Dex. Enhancers selected on differential H3K27ac-levels (DESeq2 padj < 0.05) during standard conditions. (**C**) Density scatter plot of H3K27ac ChIP-seq tag counts in hBM-MSC-TERT4 cells on day 1 of osteoblast (upper panel) or adipocyte differentiation (lower panel) with 10 and 100 nM Dex. Enhancers selected on differential H3K27ac-levels (DESeq2 padj < 0.05) during standard conditions. (**D**) Heatmap quantifying the enrichment of Dex-sensitive genes over non-regulated genes near Dex-sensitive enhancers within a window of 50 kb from the TSS. Enhancers and genes with changing activity (DESeq2 padj < 0.01) during 1 day of osteoblast (left panel) or adipocyte differentiation (right panel) were grouped into equally sized octiles based on the ranked difference in H3K27-acetylation (vertical box plots) or gene expression (horizontal box plots) when differentiating the cells with or without Dex. Number of genes or unique enhancer-gene pairs per octile depicted next to box plots (band, mean; box, first and third quartiles; whiskers, 1.5 times the interquartile range). (**E**) ROC curve for GR binding (ChIP-seq tag count) on day 1 of osteoblast (blue) or adipocyte differentiation (red) to predict increase (full) or loss (dashed) of H3K27ac signal (DESeq2 p-adj < 0.01). Insert quantifying area under the curve with p-value from roc.test function of pROC package in R. (**F**) Heat map showing H3K27-acetylation (left panel) and GR ChIP-seq tag density (right panel) within a window of +/-3 kb from the peak center. Enhancers with differential H3K27ac levels (p-adj < 0.01) during 1 day of osteoblast or adipocyte differentiation in hBM-MSC-TERT4 cells were grouped into sites with osteoblast-specific, adipocyte-specific, or common gain or loss of signal, and ordered based on induction levels. (**G**) Box plot (band, median; box, first and third quartiles; whiskers, 1.5 times the interquartile range) quantifying H3K27ac ChIP-seq tag count in hBM-MSC-TERT4 cells prior to and after 1 day of osteogenic or adipogenic differentiation with the indicated concentrations of dexamethasone. Enhancers are based on clustering in Fig. 3F with osteoblast-specific (left panel), adipocyte-specific (middle panel) and commonly upregulated H3K27ac levels (right panel). Effect sizes as Cohen’s d. (**H**) Box plot (band, median; box, first and third quartiles; whiskers, 1.5 times the interquartile range) as in Fig. 3G for GR ChIP-seq. Effect sizes as Cohen’s d. (**I**) Box plot (band, median; box, first and third quartiles; whiskers, 1.5 times the interquartile range) quantifying DNase-seq (left panel) and MED1 ChIP-seq tag count (right panel) in undifferentiated hBM-MSC-TERT4 cells at enhancers with increased H3K27ac levels according to clustering in Fig. 3F. Effect sizes ad Cohen’s d. (**J**) Box plot (band, median; box, first and third quartiles; whiskers, 1.5 times the interquartile range) quantifying the HOMER-derived log-odds score for the GC response element in genomic sequences at enhancers with increased H3K27ac levels according to clustering in Fig. 3F. Effect sizes as Cohen’s d. (**K**) Scatter plot quantifying the differentiation (circles) and Dex-dependent difference (triangles) in H3K27ac ChIP-seq levels on day 1 of osteoblast (blue) or adipocyte (red) differentiation at enhancer sites with increased H3K27ac levels according to clustering in Fig. 3F. For each group of osteoblast-specific (left panel), adipocyte-specific (middle panel) and commonly induced enhancers (right panel), enhancers were binned by GR ChIP-seq tag count on day 1 of differentiation (1d Ob 10 nM Dex and 1d Ad 100 nM Dex). Dots represent mean and error bars depict standard error. Heat map shows number of enhancers in each bin. (**L**) Model figure illustrating the binding of GR and activation of lineage-selective and commonly activated enhancers during early osteoblast and adipocyte differentiation.

### GR licenses activation of pre-established osteoblast enhancers

Next, we wanted to understand whether chromatin and sequence features as well as hormone-sensitivity and GR binding could demarcate lineage-selectivity of early enhancer activation. While enhancers with osteoblast-specific activation patterns showed higher accessibility and activity levels prior to differentiation (Fig. 3I), GRE (Glucocorticoid Responsive Element) motif strength was weaker compared to sites with adipocyte-specific or common activity profiles (Fig. 3J). Thus, pre-established chromatin accessibility at osteogenic enhancers could explain the comparable recruitment of GR under osteogenic and adipogenic conditions, while the high GRE motif score of commonly induced enhancers would fit a GR-based lineage-independent remodelling activity.

Given, the different binding patterns of GR to induced enhancers, we asked whether the magnitude of hormone-dependent enhancer activation scales with GR binding intensity under osteogenic and adipogenic conditions. Due to the lineage and hormone dose effect on GR occupancy, direct site-by-site comparisons are not informative. We therefore pooled enhancers into bins with comparable GR binding intensities and quantified lineage- and Dex-dependent changes in H3K27 acetylation across these groups. This analysis revealed a GR binding-intensity-dependent increase in H3K27 acetylation that was restricted to lineage-appropriate enhancer classes (Fig. 3K). In addition, hormone-dependent enhancer activity was almost equivalent to that of osteogenic differentiation for all groups, while adipogenic- and hormone-induced enhancer activity showed opposing patterns (Fig. 3K). The latter ranges from partial hormone-dependency at adipocyte-selective enhancers, over full at commonly activated ones, to an inverse relation at osteoblast-selective enhancers. Taken together, these findings indicate that GR occupancy provides a quantitative, dose-tuneable signal that enables enhancer activation, whereas lineage-specific chromatin features, enhancer identity, and interaction with other transcription factors dictate the selectivity and amplitude of hormone-dependent H3K27 acetylation. This outlines a model (Fig. 3L) in which commonly activated enhancers purely dependent on GR binding intensity and the presence of GRE-sequence features to drive lineage-independent remodelling. In contrast, adipocyte-selective enhancer activation is facilitated by GR binding but requires additional lineage-context likely to overcome chromatin remodelling at closed enhancers with low GRE-motif features. Lastly, the accessibility of osteogenic enhancers allows lineage-independent recruitment of GR but hormone- and GR-binding intensity dependent activation under osteogenic conditions only.

### Transcription factor MZF1 identified as an endogenous inhibitor of osteoblast differentiation

Given that GR binding alone could not explain lineage-selective enhancer activation, we next sought to identify transcription factors whose activity may modulate GC-dependent enhancer responses during OB differentiation. We applied the machine learning tool IMAGE (Madsen et al., 2017) to our H3K27ac ChIP-seq and RNA-seq data and found the expected Dex (both cocktails), vitamin D (osteogenic), and cAMP (IBMX - adipogenic) dependent motif activities of the GC and vitamin D receptor as well as the cAMP response element binding protein, respectively (Fig. 4A). In line, the strong effect of Dex on both osteoblast and adipocyte differentiation, we found a bigger repertoire of motifs changing activity upon addition of Dex to the differentiation cocktail (Supplementary Fig. 4A). Interestingly, most of these changes cannot be attributed to the transcriptional regulation of the respective transcription factors (Supplementary Figure 4B). IMAGE also revealed a stronger association between hormone- and lineage-dependent changes in motif activity upon osteogenic compared to adipogenic differentiation (Fig. 4B) indicating a strong interaction between GC-signalling and activity of other transcription factors during in osteoblast differentiation. Combining lineage-specific and hormone-specific changes in motif activity for osteoblast and adipocyte differentiation (Fig. 4C), we found two motifs, namely myeloid zinc finger 1 and zinc finger 433 (Fig. 4D), with osteoblast selective motif properties that are shaped by the presence of GC signalling. Using small interference RNA and stable expression of shRNA in hBM-MSC-TERT4 cells, we were able to reduce MZF1 (Fig. 4E & Supplementary Fig. 4C) but not ZNF433 protein levels (data not shown). Upon ablation of MZF1, cells showed higher osteogenic potential *in vitro*, as evidenced by ALP staining and mineralization (Fig. 4F & Supplementary Fig. 4D) as well as marker gene expression (Fig. 4G & Supplementary Fig. 4E), and improved bone forming capacity when implanted into immunodeficient mice (Fig. 4H). Interestingly, and not expected from the motif activity profile, loss of MZF1 increased lipid accumulation (Supplementary Fig. 4F, G, I & J) and marker gene expression (Supplementary Fig. 4H & K) upon adipogenic differentiation. Collectively, the GC- and lineage-dependent histone acetylation dynamics identified *MZF1* as an endogenous inhibitor of osteoblast differentiation in human cells.

**Figure 4.**
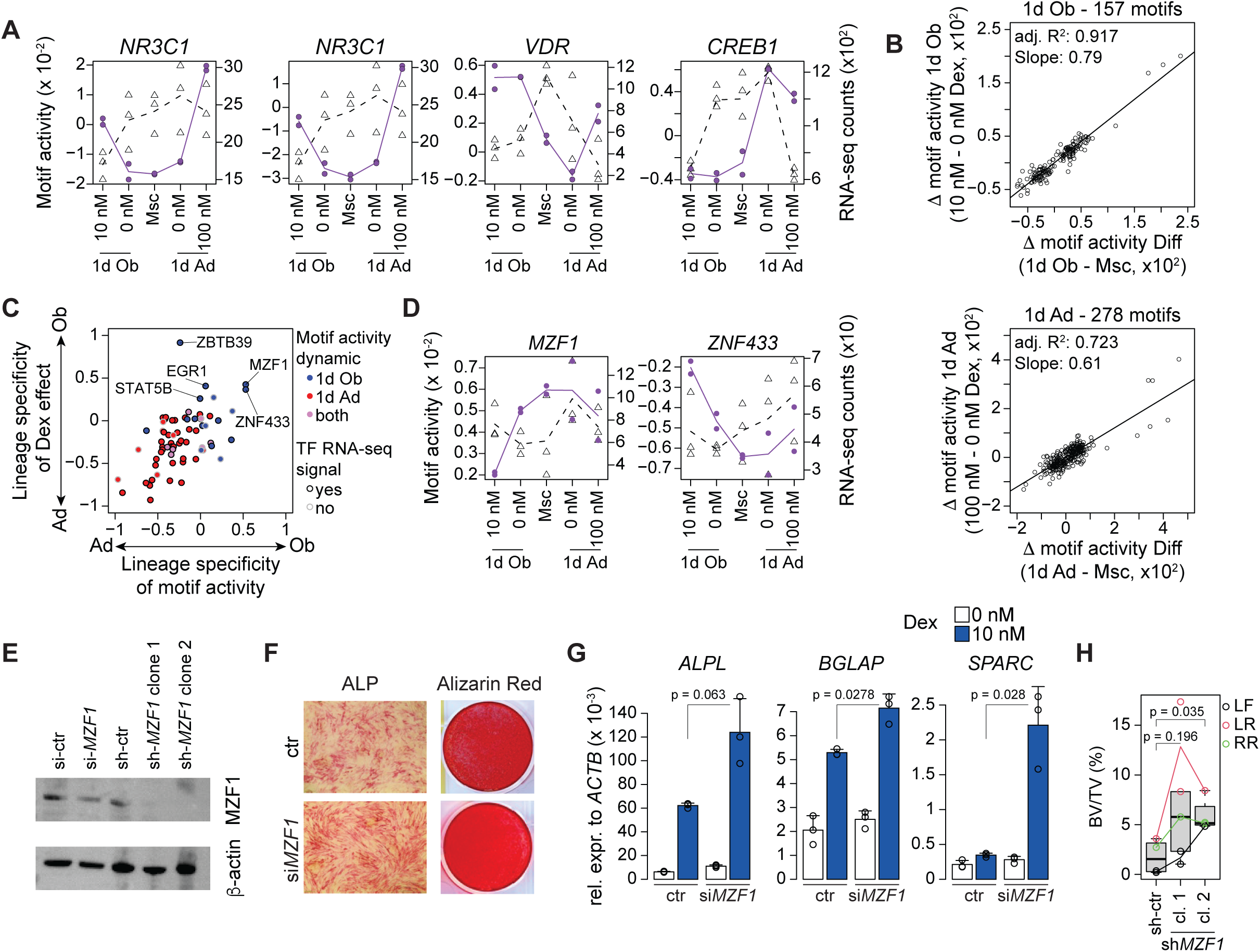
Glucocorticoid-dependent motif activity profiles identify MZF1 as endogenous inhibitor of osteoblast differentiation. (**A**) IMAGE derived motif activity profiles of selected transcription factors GR (*NR3C1* - with two motifs), vitamin D receptor (*VDR*), and cAMP response element binding protein 1 (*CREB1*) based on H3K27ac ChIP-seq in undifferentiated hBM-MSC-TERT4 cells as well as after 1 day of osteogenic and adipogenic differentiation with the indicated concentrations of Dex. Dashed lines show gene expression levels of the transcription factor (TF). (**B**) Scatter plot contrasting the differentiation and Dex-dependent changes in IMAGE-based motif activity on day 1 of osteoblast (upper panel) and adipocyte differentiation (lower panel) of hBM-MSC-TERT4 cells. Line based on linear regression with residuals. Analysis is restricted to motifs with differential activity (p-value < 0.05) for at least one condition. (**C**) Scatter plot showing lineage specificity of differentiation (delta values of x axes in Fig. 2B) and Dex-dependent (delta values of y axes in Fig. 2B) changes in IMAGE-based motif activity. Analysis is restricted to motifs with differential activity (p-value < 0.05) dependent on differentiation or Dex exposure during 1 day of osteoblast (blue), adipocyte differentiation (red), or both (purple). Black and grey outlines indicate presence or absence of RNA expression of the respective TF. (**D**) IMAGE derived motif activity profiles of myeloid zinc finger 1 (*MZF1*) and zinc finger protein 433 (*ZNF433*) based on H3K27ac ChIP-seq in undifferentiated hBM-MSC-TERT4 cells as well as after 1 day of osteogenic and adipogenic differentiation with the indicated concentrations of Dex. Dashed lines show gene expression levels of the TF. (**E**) Western blot analysis of MZF1 and β-actin in hBM-MSC-TERT4 cells with *MZF1* or control siRNA knockdown or hBM-MSC-TERT4 cells with stable expression of *MZF1* targeting shRNAs. (**F**) ALP (day 7) and Alizarin Red staining (day 14) in osteogenic differentiated hBM-MSC-TERT4 cells. Cells were treated with control or *MZF1* targeting siRNA 3 days prior to induction of differentiation. (**G**) qPCR-based mRNA expression levels of *ALPL*, *BGLAP*, and *SPARC* in hBM-MSC-TERT4 cells after 7 days of osteogenic differentiation in the presence or absence of Dex. Cells were treated with control or *MZF1* targeting siRNA 3 days prior to induction of differentiation. Bar showing mean and error bars depicting standard deviation, n = 3. Unpaired Student’s t-test. (**H**) Box plot (band, median; box, first and third quartiles; whiskers, 1.5 times the interquartile range) showing histology-based quantification of bone formed by hBM-MSC-TERT4 cells stably carrying control or shRNA constructs targeting *MZF1* after 8 weeks of ectopic transplantation into immune-deficient mice. Three different incisions at left front (LF), left rear (LR), and right rear (RR) were done on the back of the mouse. Paired t-test based on implant location.

### GC impair late maturation of human osteoblast differentiation

Since GC dosage did not substantially affect early osteogenic genes or enhancer regulation, but impaired mineralization *in vitro*, we mapped the transcriptional divergence of hBM-MSC-TERT under 10 and 100 nM Dex throughout osteoblast differentiation (Fig. 5A). The number of genes with GC-concentration-sensitive expression progressively increased with maturation (Supplementary Fig. 5A), including genes related to extracellular matrix, signalling, and metabolism (Fig. 5B) which highlighted two overall trends. First, dose-dependent regulation was poorly established on day 1 but became directionally apparent by day 7 and was further amplified on day 12 (Fig. 5C & D and Supplementary Fig. 5B & C). Second, genes induced by high-dose GC were predominantly repressed during osteoblast maturation from day 7 to day 12 and vice versa (Fig. 5E and Supplementary Fig. 5D & E). Thus, high-dose GC exposure progressively counteracts the transcriptional changes accompanying late osteoblast maturation and maintains cells in an earlier differentiation state.

**Figure 5.**
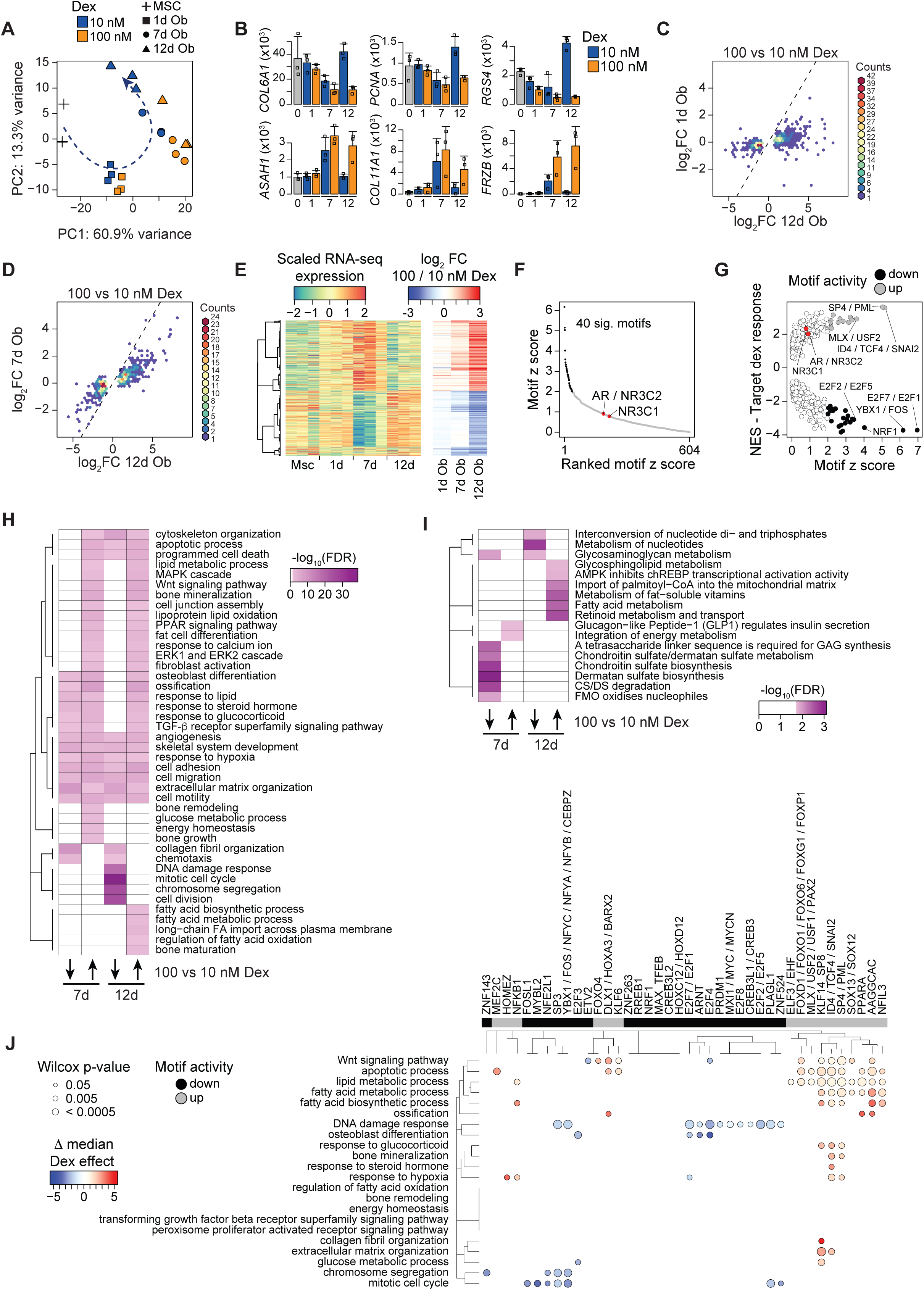
Glucocorticoid-induced inhibition of osteoblast differentiation is a cumulative late phase effect that prevents transition to a mineralizing state. (**A**) PCA plot of RNA-seq data from osteogenic stimulated hBM-MSC-TERT4 cells in the presence of 10 or 100 nM Dex. Dashed line indicates transition through standard osteoblast differentiation. (**B**) Bar plots showing RNA-seq based gene counts in hBM-MSC-TERT4 cells on days 0, 1, 4, and 12 of osteogenic differentiation with 10 or 100 nM Dex. Examples were selected based on differential expressions by Dex on day 12 (DEseq2 padj < 0.01). Bar showing mean and error bars depicting standard deviation. (**C**) Density scatter plot comparing changes in gene expression under 100 versus 10 nM Dex on days 12 and 1 of osteoblast differentiation. Genes selected based on differential expression by Dex on day 12 (DEseq2 padj < 0.01). Dashed line represents identity line. (**D**) Density scatter plot comparing changes in gene expression under 100 versus 10 nM Dex on days 12 and 7 of osteoblast differentiation. Genes selected based on differential expression by Dex on day 12 (DEseq2 padj < 0.01). Dashed line represents identity line. (**E**) Heat map showing scaled RNA-seq based expression levels throughout osteoblast differentiation for genes with differential expression by 100 vs 10 nM Dex on day 7 or 12 of osteoblast differentiation (DEseq2 padj < 0.01). Hierarchical clustering based on time course expression and side bare visualizing 100 vs 10 nM Dex effect. (**F**) Scatter plot showing the ranked z-value distribution of the ISMARA-based motif activity using RNA-seq data of hBM-MSC-TERT4 cells on day 12 of osteoblast differentiation with 10 and 100 nM of Dex. GR motifs are highlighted. (**G**) Scatter plot visualizing the z-score of the ISMARA-based motif activity from Fig. 5F and Dex response of the motif’s predicted target genes. Genes were ranked by DEseq2 Wald statistics comparing 100 vs 10 nM Dex on day 12 of osteoblast differentiation, and gene set enrichment analysis was performed for each motif using the top 500 ISMARA-predicted target genes. Motifs with significant gain and loss of activity according to Figure S5F as well as motifs for GR are highlighted. (**H**) Heat map visualizing go-seq based gene ontology enrichment (go-seq p-adj < 0.01) of genes showing differential expression by 100 versus 10 nM Dex on days 12 and 7 of osteoblast differentiation. (**I**) Heat map visualizing go-seq based REACTOME enrichment (go-seq p-adj < 0.01) of genes showing differential expression by 100 versus 10 nM Dex on days 12 and 7 of osteoblast differentiation. (**J**) Bubble plot visualizing the Dex response of motif target genes against remaining genes of selected gene ontology terms. Genes were ranked by DEseq2 Wald statistics comparing 100 vs 10 nM Dex on day 12 of osteoblast differentiation. Circle size represents a one-sided Wilcoxon rank-sum test (p-value < 0.05). Color indicates the differences in median signed Wald statistic between motif target and non-target genes. Motifs with significant gain or loss of activity according to Figure S5F are indicated.

To identify regulatory mechanisms associated with this late transcriptional arrest, we applied ISMARA to our RNA-seq data. Interestingly, GR-associated motifs showed comparatively limited differential activity between 10 and 100 nM Dex, whereas 40 motifs, some of them linked to transcription factors (TF) known to regulate cell-cycle, differentiation, and energy-sensing, displayed pronounced activity differences (Fig. 5F and Supplementary Fig. 5F). To link those TFs to the underlying biological processes we built a gene regulatory network using ISMARA-predicted target genes and found changes in motif activity to be accompanied by directional regulation of the predicted target genes (Fig. 5G), suggesting that the late high-dose response is not dominated by canonical GRE-dependent transcription but involves broader remodelling of the transcriptional network. To exclude that these associations simply reflected extensive redundancy among the predicted motif target sets, we quantified pairwise target-gene overlap and found generally low similarity (Supplementary Fig. 5G & H).

Gene ontology and Reactome pathway analyses of the GC-concentration-dependent genes identified a variety of biological programs related to cell-cycle progression, osteoblast and matrix function, lipid and fatty acid metabolism, energy homeostasis, and cellular stress (Fig. 5H, I). We therefore integrated these biological programs with the ISMARA-predicted regulatory network to identify distinct regulatory modules rather than a common GR-centered response (Fig. 5J). E2F-, MYBL2-, and NFY-associated motifs were preferentially associated with repression of mitotic and chromosome-segregation genes, whereas separate FOX-, MLX/USF-, PPARA-, and KLF-motif groups were associated with GC-sensitive genes involved in lipid and fatty-acid metabolism, energy homeostasis, stress responses, and selected osteoblast programs. Thus, high-dose GC exposure interferes with the coordinated late differentiation program at several regulatory levels instead of eliciting a uniform amplification of direct GR target-gene regulation.

### High-dose GC impairs cellular bioenergetics and induces a distinct senescence-associated stress state

The prominent involvement of energy-homeostatic and lipid-metabolic programs suggested that the impaired late transcriptional transition upon high-dose GC exposure may be accompanied by altered cellular bioenergetics. Notably, expression of energy-sensing AMPK subunits *PRKAA2* and *PRKAG2*, which normally declined from day 7 to day 12 of osteoblast differentiation, remained elevated under high-dose GC exposure on day 12 (Supplementary Fig. 6A). These findings prompted us to investigate whether high-dose GC exposure alters cellular bioenergetics during late osteoblast differentiation. Seahorse extracellular flux-based quantification of ATP production revealed substantial metabolic remodeling during osteoblast differentiation, with a progressive reduction in glycolytic ATP production and an increasing contribution of mitochondrial ATP production (Fig. 6A and Supplementary Fig. 6B). High-dose GC exposure further reduced glycolytic ATP production, resulting in lower total ATP production that was not compensated by mitochondrial ATP production (Fig. 6A & B and Supplementary Fig. 6B). Notably, the metabolic difference between 10 and 100 nM Dex was most pronounced on day 7, preceding the strongest transcriptional divergence observed on day 12. Analysis of mitochondrial respiration further revealed GC concentration-dependent differences in basal and maximal respiration as well as spare respiratory capacity (Supplementary Fig. 6C). Thus, high-dose GC exposure imposes an energetic constraint during an intermediate stage of osteoblast differentiation before manifestation of the late transcriptional maturation arrest.

**Figure 6.**
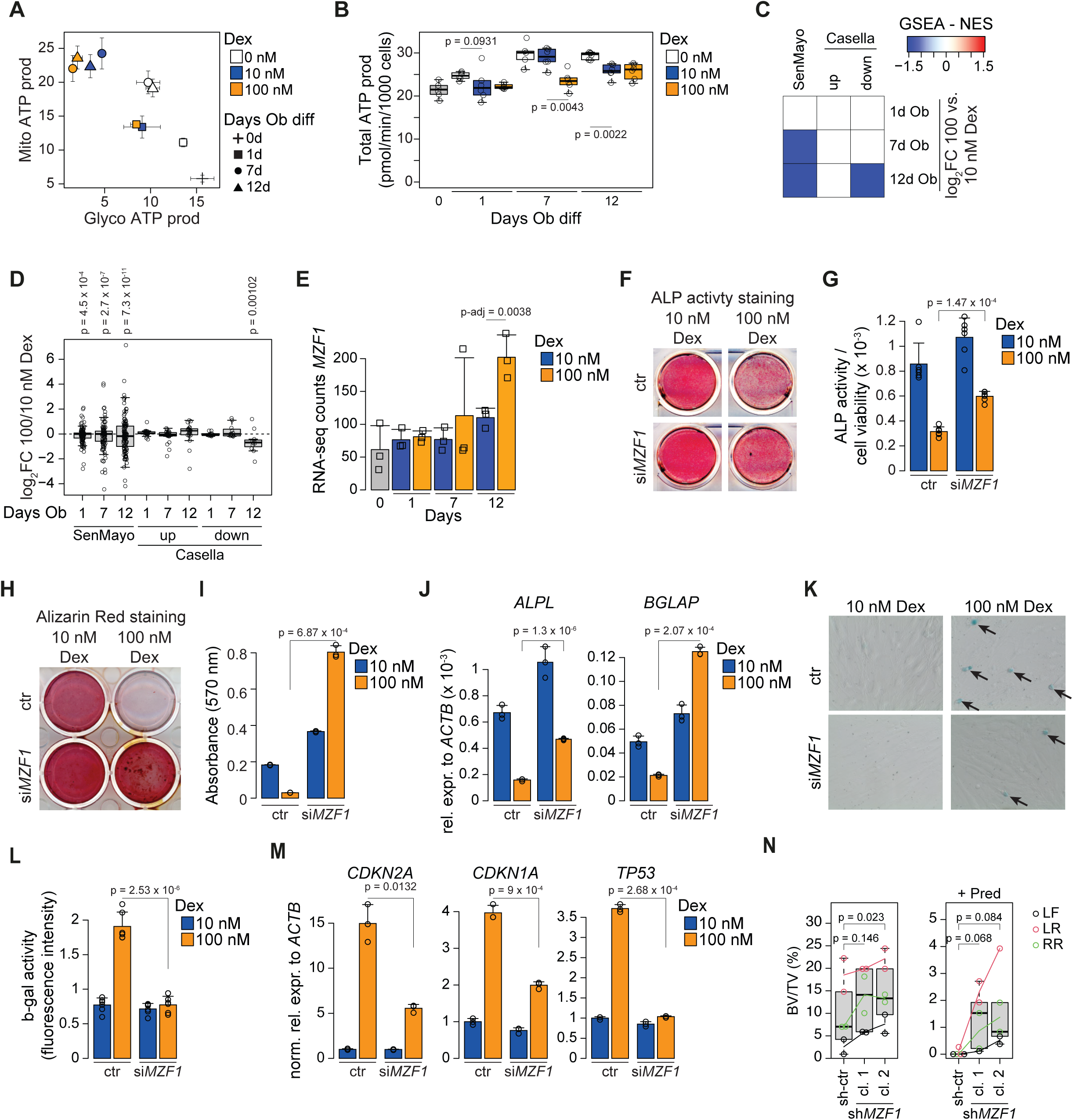
*MZF1* depletion reduces glucocorticoid-induced senescence-burden and impairment of osteoblast differentiation. (**A**) Seahorse XF analysis-based energy map for the indicated concentrations of Dex at the various time points throughout osteogenic differentiation. Mitochondrial and glycolytic ATP as means with error bars depicting standard deviation. **(B)** Box plot showing Seahorse XF analysis-based total ATP production (glycolytic + mitochondrial ATP production) for the indicated concentrations of Dex at various time points of osteoblast differentiation. (**C**) Normalized enrichment score (NES) from gene set enrichment analysis (p-adj < 0.05) based on two distinct senescence gene signatures, SenMayo (Saul et al., 2022) and a multi-trait signature from Casella et al. split into up and down regulated genes (Casella et al., 2019), and the fold changes comparing 100 vs. 10 nM Dex on days 1, 7, and 12 of osteoblast differentiation. (**D**) Box plot quantifying RNA-seq based fold changes of senescence gene signatures, Sen-Mayo (Saul et al., 2022) and up- and downregulated genes of Casella et al. (Casella et al., 2019), contrasting 100 vs. 10 nM Dex on days 1, 7, and 12 of osteoblast differentiation. P-value indicates one-tailed Wilcoxon–Mann–Whitney test comparing absolute log fold changes of signature genes against all other detected genes. (**E**) RNA-seq based counts of *MZF1* in hBMSC-TERT4-cells prior to and after 1, 7, or 12 days of osteogenic differentiation with 10 or 100 nM of Dex. DESeq2 adjusted p-value. (**F**) ALP activity in hBM-MSC-TERT4 cells on day 7 of osteogenic differentiation with 10 or 100 nM of Dex. Cells were treated with control or *MZF1* targeting siRNA 3 days prior to induction of differentiation. (**G**) Bar plot quantifying ALP-activity normalized to cell viability from cells in Fig 5B. Bar showing mean and error bars depicting standard deviation, n = 6. Unpaired Student’s t-test. (**H**) Alizarin Red staining in hBM-MSC-TERT4 cells on day 14 of osteogenic differentiation with 10 or 100 nM of Dex. Cells were treated with control or *MZF1* targeting siRNA 3 days prior to induction of differentiation. (**I**) Bar plot quantifying Alizarin Red staining intensity from cells in Fig 5E. Bar showing mean and error bars depicting standard deviation, n = 3. Unpaired Student’s t-test. (**J**) qPCR-based mRNA expression levels of *ALPL* and *BGLAP* in hBM-MSC-TERT4 cells after 7 days of osteogenic differentiation with 10 or 100 nM of Dex. Cells were treated with control or *MZF1* targeting siRNA 3 days prior to induction of differentiation. Bar showing mean and error bars depicting standard deviation, n = 3. Unpaired Student’s t-test. (**K**) Microscopic picture of β-galactosidase activity staining in hBM-MSC-TERT4 cells on day 7 of osteogenic differentiation with 10 or 100 nM of Dex. Cells were treated with control or *MZF1* targeting siRNA 3 days prior to induction of differentiation. (**L**) Fluorescence-based quantification of β-galactosidase activity in cells in Fig. 5H. Bar showing mean and error bars depicting standard deviation, n = 5. Unpaired Student’s t-test. (**M**) RT-qPCR-based mRNA expression levels of *CDKN2A* (encoding p16), *CDKN1A* (encoding p21), and *TP53* (encoding p53) in hBM-MSC-TERT4 cells after 7 days of osteogenic differentiation with 10 or 100 nM of Dex. Cells were treated with control or *MZF1* targeting siRNA 3 days prior to induction of differentiation. Bar showing mean and error bars depicting standard deviation, n = 3. Unpaired Student’s t-test. (**N**) Box plot (band, median; box, first and third quartiles; whiskers, 1.5 times the interquartile range) showing histology-based quantification of bone formed by hBM-MSC-TERT4 cells stably carrying control or shRNA constructs targeting *MZF1* after 8 weeks of ectopic transplantation into immune deficient mice. Mice underwent sham operation (left panel) or insertion of prednisolone pellets (right panel) with an approximate dose of 3 mg/kg/day. Three different incisions at left front (LF), left rear (LR), and right rear (RR) were done on the back of the mouse. Paired t-test based on implant location.

Given the association of metabolic perturbation with cellular stress and senescence (Grootaert, 2024; Song et al., 2022; Wang et al., 2023; Wiley & Campisi, 2021) on the one hand, and GC-induced osteoporosis and the emergence of senescence-associated phenotypes in bone and bone marrow (Liu et al., 2023; Poulsen et al., 2014; Wang et al., 2021; Weinstein, 2001) on the other hand, we next asked whether prolonged high-dose GC exposure was accompanied by senescence-associated transcriptional changes in our model system. We compared the GC response with the SenMayo signature, which captures senescence-associated gene expression in aged tissues including bone (Saul et al., 2022), and the Casella signature comprising genes consistently regulated across multiple experimentally induced senescence models (Casella et al., 2019). High-dose GC exposure increasingly affected these senescence-associated gene sets during differentiation, but the response differed markedly between signatures (Fig. 6C & D). Casella-down genes showed significant repression by high-dose GC, whereas Casella-up genes displayed only a weaker tendency towards induction. In contrast, the SenMayo signature shifted overall towards repression rather than the induction expected for an age-associated senescence program (Fig. 6C). Importantly, analysis of absolute fold changes demonstrated that SenMayo genes were nevertheless among the gene sets most strongly affected by high-dose GCs, indicating pronounced regulation without establishment of the age-associated senescence program captured by SenMayo (Fig. 6D). Casella-down genes were likewise significantly affected in magnitude, consistent with their coherent repression. Moreover, regulation of SenMayo and Casella-up genes by high-dose GCs opposed their regulation during the normal transition from day 7 to day 12 (Supplementary Fig. 6D).

### MZF1 sustains the GC-induced stress state and impairment of osteoblast differentiation

Given the inhibitory effect of *MZF1* on osteoblast differentiation and its reported involvement in senescence-relevant transcriptional control circuits, including regulation of p16^INK4A^ and p53-associated programs (Li et al., 2023; Wu et al., 2022), we asked whether *MZF1* expression was affected by prolonged high-dose GC exposure. Indeed, *MZF1* was upregulated hy high-dose GCs on day 12 of osteoblast differentiation (Fig. 6E), alongside induction of the p21-encoding *CDKN1A* (Supplementary Fig. 6E). We therefore questioned whether MZF1 contributes to cellular stress and the late-stage differentiation arrest induced by high-dose GC. Depletion of MZF1 attenuated the inhibitory effect of 100 nM Dex on ALP activity and matrix mineralization (Fig. 6F-I and Supplementary Fig. 6F-H) and restored expression of the osteoblast markers *ALPL* and *BGLAP* (Fig. 6J and Supplementary Fig. 6I). *MZF1* knockdown also reduced the high-dose GC-induced increase in senescence-associated β-galactosidase activity (SA-β-gal) (Fig. 6K & L) and attenuated expression of the cell-cycle and stress-associated genes *CDKN2A*, *CDKN1A*, and *TP53* (Fig. 6M and Supplementary Fig. 6J). Finally, prednisolone (Pred) reduced ectopic bone formation by control hBM-MSC-TERT cells *in vivo*, whereas MZF1-depleted cells retained greater bone-forming capacity during GC exposure (Fig. 6N). Together, these findings identify MZF1 as a component of a regulatory network that sustains the cellular stress and maturation arrest induced by prolonged high-dose GC exposure.

## Discussion

GC exert seemingly contradictory effects on skeletal cells, being required for osteoblast differentiation and skeletal homeostasis at physiological levels while suppressing bone formation upon prolonged exposure or at pharmacological concentrations (Chen et al., 2023; Gado et al., 2022; Martin et al., 2021). Our findings suggest that these opposing actions are not simply different points along a linear GR dose-response but instead reflect distinct context- and concentration-dependent modes of GC action. Cellular context determines the immediate regulatory consequences of GR activation, whereas the adverse effects of GC excess develop over time and cannot be predicted from the early transcriptional response.

This distinction provides a framework for reconciling the permissive and inhibitory effects of GC on osteoblast differentiation. Importantly, our *in vitro* system of the hBM-MSC-TERT4 cells does not aim to recapitulate the complex pathophysiology of GC-induced osteoporosis, which integrates systemic and cell-type-specific effects across multiple skeletal and non-skeletal tissues (Chotiyarnwong & McCloskey, 2020; Hofbauer et al., 2025). Instead, it allows receptor dependence, lineage context, GC concentration and differentiation stage to be experimentally separated within the same human cellular background. Our data thereby identify two conceptually distinct layers of GC action: an early GR-dependent licensing of lineage-specific regulatory programs and a cumulative response to GC excess that interferes with late osteoblast maturation.

### GR binding licenses lineage-specific enhancer activation in a context-dependent manner

GR recruitment was closely linked with enhancer activation during early stromal cell differentiation, supporting a direct regulatory contribution of GR to the establishment of lineage-specific transcriptional programs. Increasing GR occupancy was accompanied by increasing enhancer activity with a particularly direct relationship during osteoblast differentiation, where GC-dependent changes in H3K27ac largely reflected the contribution of GR. However, this relationship did not fully explain where enhancer activation occurred. GR occupancy increased with GC concentration across enhancer classes and, importantly, could occur at lineage-selective enhancers outside the cellular context in which these enhancers became activated. Thus, GR binding is required and quantitatively linked to enhancer activation, but its regulatory consequence is conditional on the surrounding lineage context.

The regulatory consequence of transcription factor binding is increasingly understood to depend on the surrounding enhancer context rather than occupancy alone. This principle has also been observed for GR: despite widespread receptor binding, only a subset of GR-bound regions possesses autonomous GC-responsive enhancer activity (Vockley et al., 2016), while substantial chromatin binding of a functionally impaired GR mutant is accompanied by markedly reduced enhancer remodeling and transcriptional output (Johnson et al., 2021). A similar dissociation between receptor occupancy and regulatory output has been observed for other ligand-activated nuclear receptors (Hah et al., 2013; Huang et al., 2021). More recent work further indicates that GR occupancy and H3K27ac are generally closely coupled, but that the strength and dynamics of this relationship depend on GRE strength, chromatin accessibility and the recruitment of additional TF (Stavreva et al., 2026). This is consistent with emerging experimental models of enhancer regulation in which combinations of context-dependent TF determine whether and to what extent the binding of an individual factor becomes transcriptionally productive (Kribelbauer-Swietek et al., 2024).

Our data suggests that the relative contribution of these determinants differs between enhancer classes. Common GC-responsive enhancers contained stronger GREs and showed a close quantitative relationship between GR occupancy and enhancer activity, consistent with a comparatively direct GC-signalling-driven response. Osteoblast-selective enhancers, in contrast, were characterized by greater pre-existing accessibility and weaker GRE. At these sites, GR could bind under both differentiation conditions, but productive GC-dependent activation was restricted to the osteogenic context. Adipocyte-selective enhancers represented yet another configuration, in which GR binding remained required but was embedded within the extensive enhancer remodeling accompanying adipogenesis in those cells (Madsen et al., 2020; Rauch et al., 2019). This is consistent with the cooperative recruitment of GR and adipogenic transcription factors such as C/EBPβ during establishment of the adipocyte enhancer landscape (Siersbaek et al., 2017; Siersbaek et al., 2011; Siersbaek et al., 2014; Steger et al., 2010).

Together, these findings argue for a model in which GR provides a ligand-dependent licensing signal rather than independently specifying enhancer identity. The amount of GR recruited contributes quantitatively to enhancer activation, while the surrounding chromatin architecture and differentiation-dependent transcription factor network determine how this input is translated into lineage-specific activity. The identification of *MZF1* as a GC-sensitive component of the osteogenic regulatory network provides one example of such contextual regulation. While *MZF1* has previously been implicated in the transcriptional regulation of *CDH2* (N-cadherin) in human osteoblasts (Le Mee et al., 2005) and *TGFB1* in human stromal cells (Weber et al., 2015), its role in physiological osteoblast differentiation has received little attention. Notably, depletion of MZF1 has been shown to promote osteoblast maturation of osteosarcoma stem-like cells through regulation of *YAP1* expression (Verma et al., 2017). Our findings extend these observations to non-transformed human stromal cells and demonstrate that *MZF1* restrains both osteoblast differentiation and bone-forming capacity.

### GC excess progressively arrests late osteoblast maturation

The consequences of GC concentration became increasingly apparent as osteoblast differentiation progressed. Importantly, the high-dose response increasingly opposed the transcriptional changes accompanying normal late osteoblast maturation *in vitro*, resulting in a state resembling an intermediate stage of differentiation.

Dose- and time-dependent effects of GC on human osteoblast differentiation *in vitro* have been previously observed at both the phenotypic as well as transcriptional level. For the former, the timing of GC treatment appears crucial, with high GC concentrations being strongly osteogenic when applied during the first week of differentiation (Alm et al., 2012). At the transcriptional level, increasing GC concentrations induced chondrogenic and adipogenic master regulators, namely *SOX9* and *PPARG*, in mature osteoblasts (Della Bella et al., 2021), as well as WNT-signalling components such as *DKK1* (Ohnaka et al., 2004).

Much of the mechanistic evidence for GC-mediated inhibition of osteogenesis, however, derives from experimental models in which pharmacological GC exposure is compared with untreated controls. Particularly in murine osteoblast systems, these approaches have identified GC-sensitive pathways whose restoration can counteract impaired differentiation, including IL-11, BMP and WNT signalling, CDK5 and cGMP/PKG signalling, as well as more recently altered Basigin signalling (Ambrosi et al., 2025; Hildebrandt et al., 2018; Kruger et al., 2022; Luppen et al., 2003; Pal China et al., 2024; Rauch et al., 2010). Similar observations have been made in human osteoblasts, where Dex reduced β-catenin/TCF activity and silencing of the WNT antagonist *DKK1* partially rescued osteoblast differentiation (Butler et al., 2010). Importantly, however, the effects of WNT signalling itself depend on the stage of human osteoblast differentiation, promoting osteogenic competence at earlier stages while inhibiting matrix mineralization when activated during later differentiation (Eijken et al., 2008). Thus, pathways identified through rescue of GC-mediated inhibition are not necessarily uniformly pro-osteogenic throughout differentiation.

This distinction is particularly relevant because human bone marrow stromal cell models have long demonstrated that GC exposure itself is required during an early window of osteoblast differentiation (Cheng et al., 1994; Eijken et al., 2006). Consequently, experiments comparing pharmacological GC exposure with untreated cells address a different question from experiments distinguishing permissive GC signalling from GC excess. In our system, increasing Dex from a permissive 10 nM to 100 nM produced comparatively limited transcriptional differences early in differentiation, whereas the major divergence accumulated after establishment of the osteogenic program. Our data therefore extends previous observations of dose-dependent regulation of individual GC-responsive genes to the broader transcriptional program and suggests that GC excess does not simply suppress osteoblast identity but increasingly interferes with progression toward a mature osteoblast state (Chen et al., 2008; Pattappa et al., 2011).

### Metabolic constraint accompanies an MZF1-dependent cellular response to GC excess

Osteoblast differentiation requires substantial metabolic adaptation, as human stromal cells increase mitochondrial activity and oxidative phosphorylation during osteogenic differentiation, and interference with this metabolic transition impairs osteoblast differentiation (Chen et al., 2008; Guo et al., 2015; Pattappa et al., 2011). This is supported by the need of mitochondrial fatty-acid oxidation in mature osteoblasts for murine bone acquisition (Alekos et al., 2023; Kim et al., 2017). Notably, oxidative metabolism appears particularly important during intermediate stages of osteoblast differentiation rather than representing a simple metabolic endpoint of maturation (Sabini et al., 2023). In our cells, the metabolic effect of GC concentration was likewise most pronounced on day 7, when high-dose GC further reduced glycolytic ATP production without compensatory maintenance of total ATP output. Previous studies have described time-dependent effects of GCs on osteoblast mitochondrial function, with prolonged exposure associated with oxidative stress and mitochondrial dysfunction (Hsu et al., 2020). Together with the temporal precedence of the metabolic phenotype over the strongest transcriptional divergence in our system, these observations raise the possibility that GC excess compromises the metabolic adaptation required for progression through an intermediate stage of osteoblast maturation.

In parallel with these metabolic changes, prolonged GC excess induced several features associated with cellular senescence. Direct evidence for GC-induced senescence in human skeletal cells remains limited, although Dex has been shown to increase SA-β-gal activity in primary human osteoblasts and other musculoskeletal cell types (Chen et al., 2021; Poulsen et al., 2014). More extensive evidence comes from murine models, where prolonged GC exposure induces senescence in skeletal stromal, adipocyte-lineage and osteoblastic cells, and genetic or pharmacological interference with these responses attenuates impaired bone formation and bone loss (Cao et al., 2023; Liu et al., 2023; Wang et al., 2021). Our data support the occurrence of a senescence-associated response also in human osteoblast differentiation but argue against interpreting this state as canonical ageing-associated senescence. Although genes of the SenMayo signature (Saul et al., 2022) were strongly affected by GC excess, their overall regulation was slightly decreased opposite to that associated with ageing, while the broader Casella signature (Casella et al., 2019) showed only partial concordance. Thus, the combination of senescence-associated β-galactosidase activity and cell-cycle regulatory changes in our cells appears to reflect a senescence-associated cellular stress state rather than accelerated cellular ageing.

*MZF1* emerged as a functional component of this prolonged response to GC excess. While MZF1 constrained osteoblast differentiation already under permissive GC conditions, prolonged high-dose GC exposure further increased *MZF1* expression together with *CDKN1A*. Importantly, depletion of MZF1 attenuated the inhibitory effect of high-dose GC on osteoblast differentiation, reduced SA-β-gal activity and expression of *CDKN2A*, *CDKN1A* and *TP53*, and preserved greater bone-forming capacity during prednisolone exposure. These findings are consistent with previous evidence linking *MZF1* to transcriptional control of *CDKN2A/p16INK4A* during oncogene-induced senescence (Wu et al., 2022), modulation of p53 during cell cycle arrest (Alekos et al., 2023), regulation of glycolytic genes such as *GAPDH*, *HK2*, and *PGK1* (Fang et al., 2020; Liu et al., 2026; Piszczatowski et al., 2014), and energy sensing AMPK signalling (Li et al., 2024). These observations place MZF1 at a potential intersection between metabolic adaptation and senescence and cell-cycle -associated transcriptional control. While our data does not establish whether the metabolic changes induced by GC excess contribute to MZF1 activation, they identify MZF1 as a functional component sustaining the differentiation-inhibitory and stress-associated state that develops during prolonged GC exposure.

### GC exposure defines distinct experimental and biological states

Our findings emphasize that the cellular consequences of GC exposure cannot be interpreted from concentration or GR occupancy in isolation. In human osteoblast differentiation, the absence of GC signalling, permissive GR activation and prolonged GC excess represent biologically distinct conditions. Their consequences depend on differentiation state and exposure duration, ranging from context-dependent enhancer activation early in differentiation to metabolic and stress-associated changes accompanying impaired maturation during prolonged hi-dose GC exposure.

This distinction is particularly important when interpreting mechanisms of GC-mediated inhibition identified by comparison with GC-naïve cells. A pathway that differs between GC-treated and untreated cells may contribute to the permissive actions required for differentiation, the detrimental consequences of GC excess, or both. Experimentally separating these states, as done here within the same human cellular background, therefore provides a way to distinguish mechanisms associated with GC signalling itself from those specifically emerging with GC excess. Such a distinction will be important for determining which cellular mechanisms identified in experimental models are relevant to the detrimental skeletal consequences of prolonged GC exposure in humans.

## Ethics declaration

Ethical approval for animal experiments was given by the Animal Experimentation Council of the Ministry of Food, Agriculture and Fishing Denmark (License 2022-15-0201-01225).

## Data availability

Databases and links will be available upon publication. Raw sequencing data reported in this study have been deposited under the NCBI Gene Expression Omnibus: GSEXXXX. Processed data and scripts for data processing and visualization to recapitulate the analyses are available at XXX and at GitHub: https://github.com/drarauch/GR_GC_Osteoblast.

## Acknowledgements

This work was supported by a grant from the Lundbeck Foundation (R335-2019-2195) and Novo Nordisk Foundation (NNF22OC0078257) to A.R. We thank Tina Nielsen for help with cell culture, Paula Fernandez-Guerra and Camilla Poulsen for assistance with Seahorse XF assays. Sequencing was carried out at the Villum Center for Bioanalytical Sciences, Functional Genomics & Metabolism Research Unit, University of Southern Denmark. We thank Ronni Nielsen for sequencing assistance.

## Author information

M.R.M., D.A., and M.P. contributed to the planning and execution of the experimental work, analyzing data, and writing the manuscript. A.C. performed knockdown and implantation experiments. V.A. made cell lines with stable shRNA expressions. A.K.H. assisted with bioinformatic analysis and training. B.B., M.K., and S.M. provided critical supervision and infrastructure in all phases of the study. A.R. was the principal supervisor, involved in planning and designing the study, analyzing data, and writing the manuscript. All authors provided critical feedback during the experimental phase and were involved in reviewing the manuscript.

