## Supplementary for "Glucocorticoid signalling exerts distinct lineage-permissive and dose-dependent disruptive actions during human osteoblast differentiation"

### Supplementary table - reagents

| Antibodies | Source | Identifier |
|---|---|---|
| anti-b-Actin | Sigma-Aldrich | A3853 |
| anti-GR | Santa Cruz | sc-1003 |
| anti-H3K27ac | Abcam | ab4729 |
| anti-Mouse IgG-HRP conjugate | Cell Signaling | #7076 |
| anti-MZF1 | Novus Biologicals | NBP2-93459 |
| anti-Rabbit IgG HRP conjugate | Cell Signaling | #7074 |
| anti-TF2B | Santa Cruz | sc-225 |
| **Oligonucleotides** | **Source** | **Identifier** |
| real time PCR primer - *ACTB* | Pentabase | For: ATTGGCAATGAGCGGTTCCG<br>Rev: AGGGCAGTGATCTCCTTCTG |
| real time PCR primer - *ADIPOQ* | Pentabase | For: GGGCCCCAGGCCGTGATGGCA<br>Rev: CGGGGACCTTCAGCCCCGGGTA |
| real time PCR primer - *ALPL* | Pentabase | For: ACGTGGCTAAGAATGTCATC<br>Rev: CTGGTAGGCGATGTCCTTA |
| real time PCR primer - *BGLAP* | Pentabase | For: CATGAGAGCCCTCACA<br>Rev: AGAGCGACACCCTAGAC |
| real time PCR primer - *CDKN1A* | Pentabase | For: GGCAGACCAGCATGACAGATT<br>Rev: GGCTTCCTCTTGGAGAAGATCA |
| real time PCR primer - *CDKN2A* | Pentabase | For: GGGTCGGGTAGAGGAGGTG<br>Rev: CATCATGACCTGGATCGGC |
| real time PCR primer - *CEBPA* | Pentabase | For: AACCTTGTGCCTTGGAAATG<br>Rev: CTGTAGCCTCGGGAAGGAG |
| real time PCR primer - *MZF1* | Pentabase | For: CTGAGGAGGCCCAGAGATGT<br>Rev: GAACCCTGGGGAGAAGATGC |
| real time PCR primer - *PPARG2* | Pentabase | For: TTCTCCTATTGACCCAGAAAGC<br>Rev: CTCCACTTTGATTGCACTTTGG |
| real time PCR primer - *SPARC* | Pentabase | For: AGAAGGAGGAGGCCGGGGCAAGG<br>Rev: CCCGCGAATGTGGAGGGGTTTA |
| real time PCR primer - *TP53* | Pentabase | For: CCTGAGGTTGGCTCTGACTGTA<br>Rev: GTTCCGTCCCAGTAGATTACCA |
| shRNA - control **Scramble** **Linker** | IDT | TCAGTCGCGTTTGCGACTGGTTCAAGAG<br>ACCAGTCGCAAACGCGACTGATTTTTTC |
| shRNA - *MZF1* 1 **Antisense** **Linker** | IDT | TGACTCTGAGGAGGAGGGTGATTCAAGA<br>GATCACCCTCCTCCTCAGAGTCTTTTTTC |
| shRNA - *MZF1* 2 **Antisense** **Linker** | IDT | TCGTACACACGGGCGAGAAATTCAAGA<br>GATTTCTCGCCCGTGTGTACGTTTTTTC |
| siRNA - *MZF1* | ThermoFischer | 4390771 (n290339) |
| siRNA - *NR3C1* | ThermoFischer | 4390824 (s6186) |
| siRNA - scramble control | ThermoFischer | 4390846 |
| **Reagents and assays** | **Source** | **Identifier** |
| 3-isobutyl-1-methylxanthine (IBMX) | Sigma-Aldrich | I5879 |
| Acetic acid | VWR | 20302236 |
| Acetone | Sigma-Aldrich | 179124 |
| Aldosterone | Sigma-Aldrich | A9477 |
| Alizarin Red | Sigma-Aldrich | A5533 |
| Alkaline Phosphatase Detection Kit | Sigma-Aldrich | SCR004 |
| a-MEM | Gibco | 31095 |
| Ascorbic acid | Wako | #013-12061 |
| BSA | Sigma-Aldrich | A4919 |
| CellTiter-Blue | Promega | G8081 |
| Chloroform | Sigma-Aldrich | C2432 |
| Citrate buffer | Sigma-Aldrich | 168782 |
| cOmplete™ protease inhibitor cocktail | Roche | 05892970001 |
| Cortisol (Hydrocortisone) | Sigma-Aldrich | H4001 |
| Dexamethsaone | Sigma-Aldrich | D4902 |
| DMEM | Gibco | 41965-039 |
| EconoSpin® RNA/DNA Micro Spin Column | Epoch Life Science | 3010 |

| | | |
|---|---|---|
| Ethanol | Sigma-Aldrich | 51976 |
| Fast SYBR™ Green Master Mix | Applied Biosystems | 4385612 |
| Formaldehyde | Sigma-Aldrich | 47608 |
| Formic acid | Sigma Aldrich | F0507 |
| Glucose | Sigma-Aldrich | G8769 |
| Glutamine | Sigma-Aldrich | G7513 |
| Hoechst 33342 | Invitrogen | H3570 |
| Insulin | Sigma-Aldrich | I9278 |
| Isopropanol | Sigma-Aldrich | I9516 |
| Lipofectamin™ 2000 | ThermoFisher | 11668019 |
| Methanol | Sigma-Aldrich | 65548 |
| Mifepristone | Sigma-Aldrich | M8046 |
| N-GARDE Mycoplasma PCR Reagent set | BioNordika (Euroclone) | EC-EMK090020 |
| NuPAGE™ Bis-Tris Mini Protein Gels, 10% | Invitrogen | NP0315BOX |
| NuPAGE™ LDS Sample Buffer (4X) | ThermoFisher | NP0007 |
| NuPAGE™ Sample Reducing Agent (10X) | ThermoFisher | NP0009 |
| Oil Red O | Sigma-Aldrich | O0625 |
| Opti-MEM™ | ThermoFisher | 31985070 |
| PBS without $Ca2^+$ and $Mg2^+$ | Invitrogen | 14190-169 |
| Penicillin Streptomycin | Gibco | #15140-122 |
| Pierce™ BCA Protein Assay Kits | ThermoFisher | 23227 |
| Pierce™ ECL Western | ThermoFisher | 32106 |
| Placebo pellets | Innovative Research of America | SC-111 |
| p-nitrophenyl phosphate | Sigma-Aldrich | 71768 |
| Prednisolone pellets (7.5 mg – 60 day release) | Innovative Research of America | SG-151 |
| pSICO | Addgene | 11579 |
| RevertAid First Strand cDNA Synthesis Kit | ThermoFisher | K1622 |
| RNA Kit (15NT) | Agilent | DNF-471-0500 |
| Rosiglitazone | Cayman Chemicals | 71740 |
| RPE buffer | Qiagen | 1018013 |
| SDS MW ladder PageRuler | ThermoFischer | PI26630 |
| SDS MW ladder Prestained 11 - 190 kDa | New England Biolabs | #P7706 |
| Seahorse XF DMEM medium | Agilent | #103575-100 |
| Seahorse XF T Cell Metabolic Profiling Kit | Agilent | #103771-100 |
| Senescence β-Galactosidase Activity Assay Kit | Cell Signaling | #23833 |
| Senescence β-Galactosidase Staining Kit | Cell Signaling | #9860 |
| Sodium hydroxide | Sigma-Aldrich | 221465 |
| Sodium pyruvate | Gibco | 11360-039 |
| TRI Reagent® | Sigma-Aldrich | 15596018 |
| Trypsin | ThermoFischer | 25300062 |
| TWEEN® 20 | MERCK | 655205 |
| Vitamin D | Sigma-Aldrich | D1530 |
| β-glycerolphosphate | Sigma-Aldrich | 50020 |

#### Supplementary figures and legends

### Supplementary Figure 1

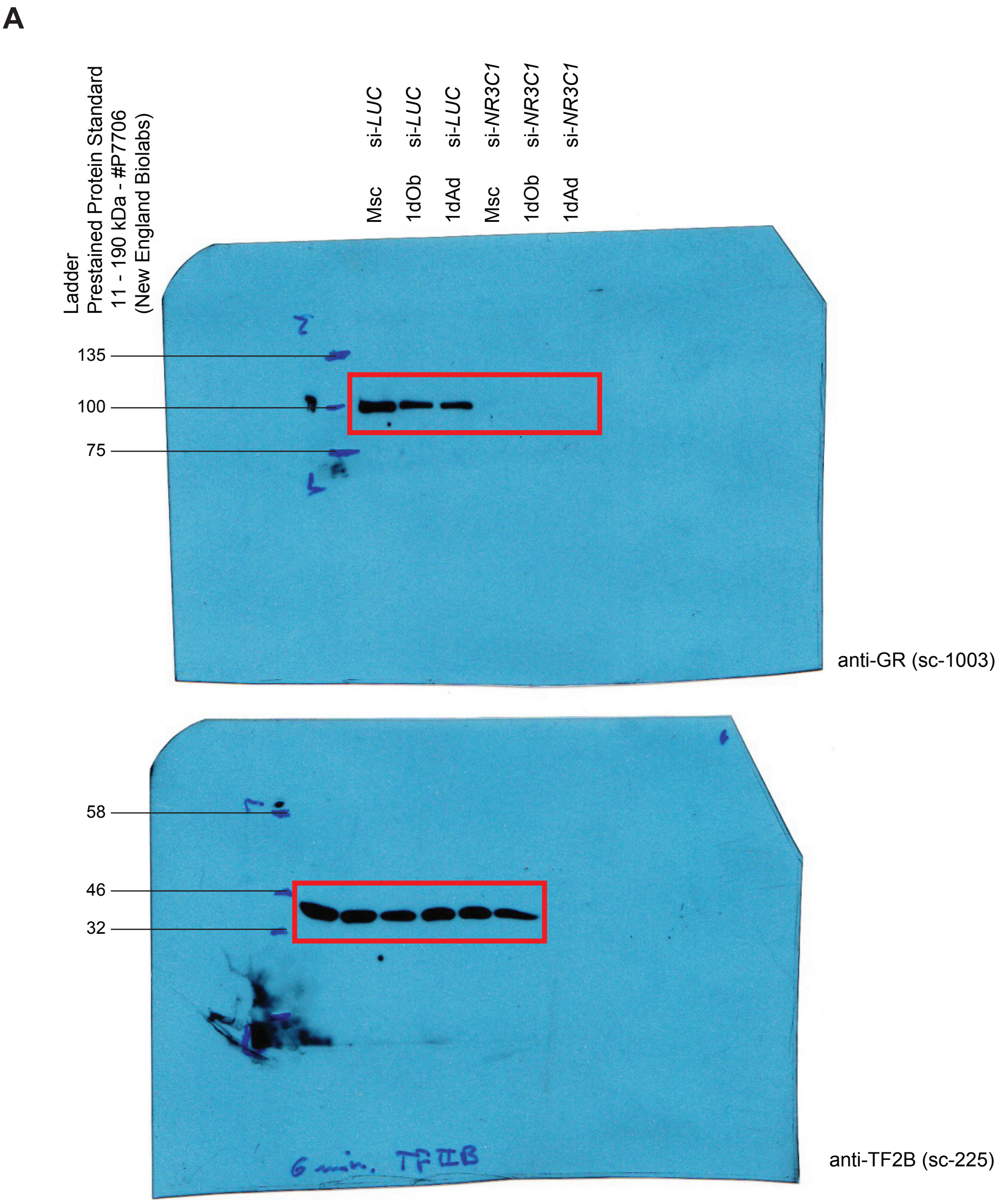

**Supplementary Figure 1 – related to Figure 1.** (**A**) Uncropped Western blot analysis shown in Figure 1E. Membrane was cut horizontally for incubation with primary antibodies against GR (top) and TF2B (bottom). Red boxes indicate the cropped region shown in Figure 1E.

### Supplementary Figure 2

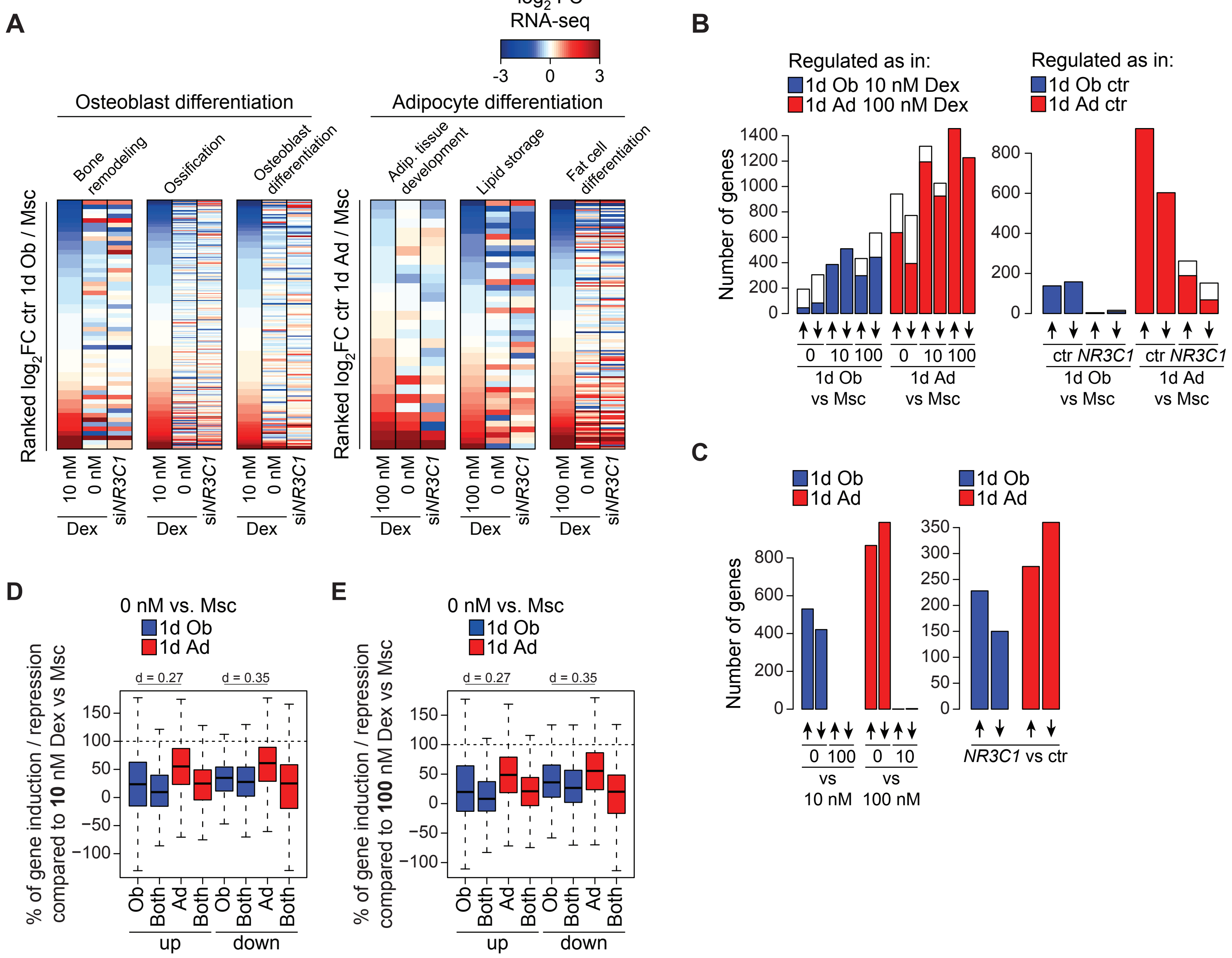

**Supplementary Figure 2 – related to Figure 2.** (**A**) Heat map showing RNA-seq based fold changes for selected gene ontology terms during 1 day of osteoblast (left panel) or adipocyte differentiation (right panel). hBM-MSC-TERT4 cells were either differentiated with or without Dex or treated with control or *NR3C1* targeting siRNA three days prior to differentiation. Fold changes in comparison to undifferentiated cells. Genes were filtered based on expression across all conditions and ranked according to fold changes upon standard conditions. (**B**) Bar plot quantifying the number of differentially expressed genes (DESeq2 adjusted p-value < 0.01) in hBM-MSC-TERT4 cells on day 1 of differentiation compared to undifferentiated cells. Cells were differentiated with indicated concentrations of Dex (left panel) or treated with control or *NR3C1* targeting siRNA 3 days prior to induction of differentiation (right panel). Colored bars indicate genes with similar regulations as under standard conditions. (**C**) Bar plot quantifying the number of differentially expressed genes (DESeq2 adjusted p-value < 0.01) in hBM-MSC-TERT4 cells on day 1 of differentiation comparing indicated concentrations of Dex (left panel) or *NR3C1* targeting versus control siRNA (right panel). (**D**) Box plot (band, median; box, first and third quartiles; whiskers, 1.5 times the interquartile range) quantifying gene regulation on day 1 of osteoblast (blue) or adipocyte (red) differentiation without Dex, normalized to the respective gene regulation with 10 nM Dex (100%). Gene groups are based on Figure 2F. Effect sizes as Cliff's delta. (**E**) Box plot (band, median; box, first and third quartiles; whiskers, 1.5 times the interquartile range) quantifying gene regulation on day 1 of osteoblast (blue) or adipocyte (red) differentiation without Dex, normalized to the respective gene regulation with 100 nM Dex (100%). Gene groups are based on Figure 2F. Effect sizes as Cliff's delta.

### Supplementary Figure 3

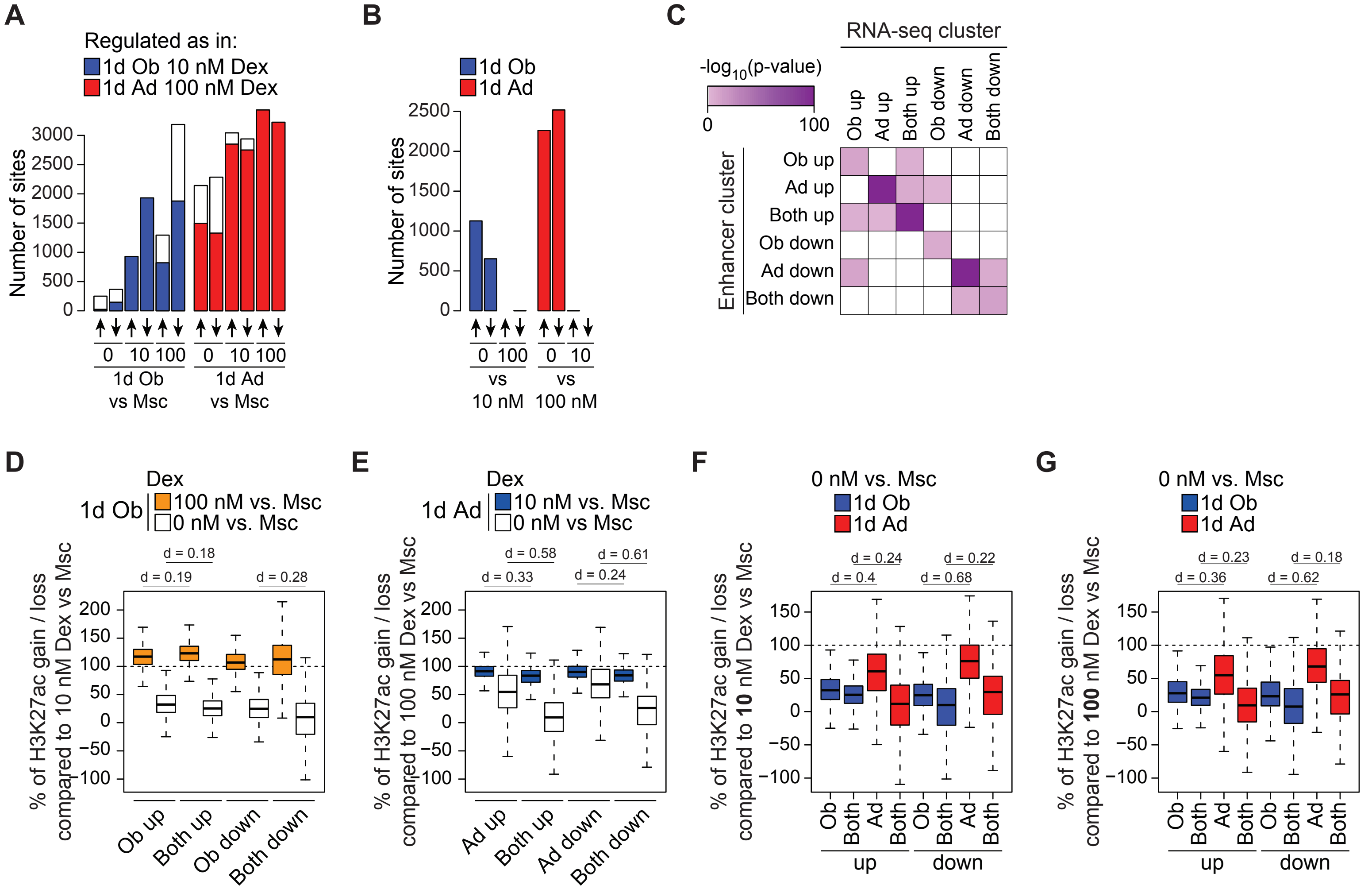

**Supplementary Figure 3 – related to Figure 3.** (**A**) Bar plot quantifying the number of enhancers with differential H3K27-acetylation (DESeq2 adjusted p-value < 0.01) in hBM-MSC-TERT4 cells on day 1 of differentiation compared to undifferentiated cells. Cells were differentiated with indicated concentrations of Dex. Colored bars indicate enhancers with similar regulations as under standard conditions. (**B**) Bar plot quantifying the number of enhancers with differential H3K27-acetylation (DESeq2 adjusted p-value < 0.01) in hBM-MSC-TERT4 cells on day 1 of differentiation comparing indicated concentrations of Dex. (**C**) Functional association of enhancer clusters with RNA-seq clusters through enrichment of human adipose tissue eQTLs ($P < 1 \times 10^{-4}$, $n = 720{,}304$) in enhancer regions. Enrichment is based on a hypergeometric test. (**D**) Box plot (band, median; box, first and third quartiles; whiskers, 1.5 times the interquartile range) quantifying changes in H3K27-acetylation levels on day 1 of osteoblast differentiation, normalized to the respective changes during standard osteoblast differentiation (100 %). Enhancer groups are based on Figure 3F. Effect sizes as Cliff's delta. (**E**) Box plot (band, median; box, first and third quartiles; whiskers, 1.5 times the interquartile range) quantifying changes in H3K27-acetylation levels on day 1 of adipocyte differentiation, normalized to the respective changes during standard adipocyte differentiation (100 %). Enhancer groups are based on Figure 3F. Effect sizes as Cliff's delta. (**F**) Box plot (band, median; box, first and third quartiles; whiskers, 1.5 times the interquartile range) quantifying changes in H3K27-acetylation levels on day 1 of osteoblast (blue) or adipocyte (red) differentiation without Dex, normalized to the respective changes with 10 nM Dex (100%). Enhancer groups are based on Figure 3F. Effect sizes as Cliff's delta. (**G**) Box plot (band, median; box, first and third quartiles; whiskers, 1.5 times the interquartile range) quantifying changes in H3K27-acetylation levels on day 1 of osteoblast (blue) or adipocyte (red) differentiation without Dex, normalized to the respective changes with 100 nM Dex (100%). Enhancer groups are based on Figure 3F. Effect sizes as Cliff's delta.

### Supplementary Figure 4

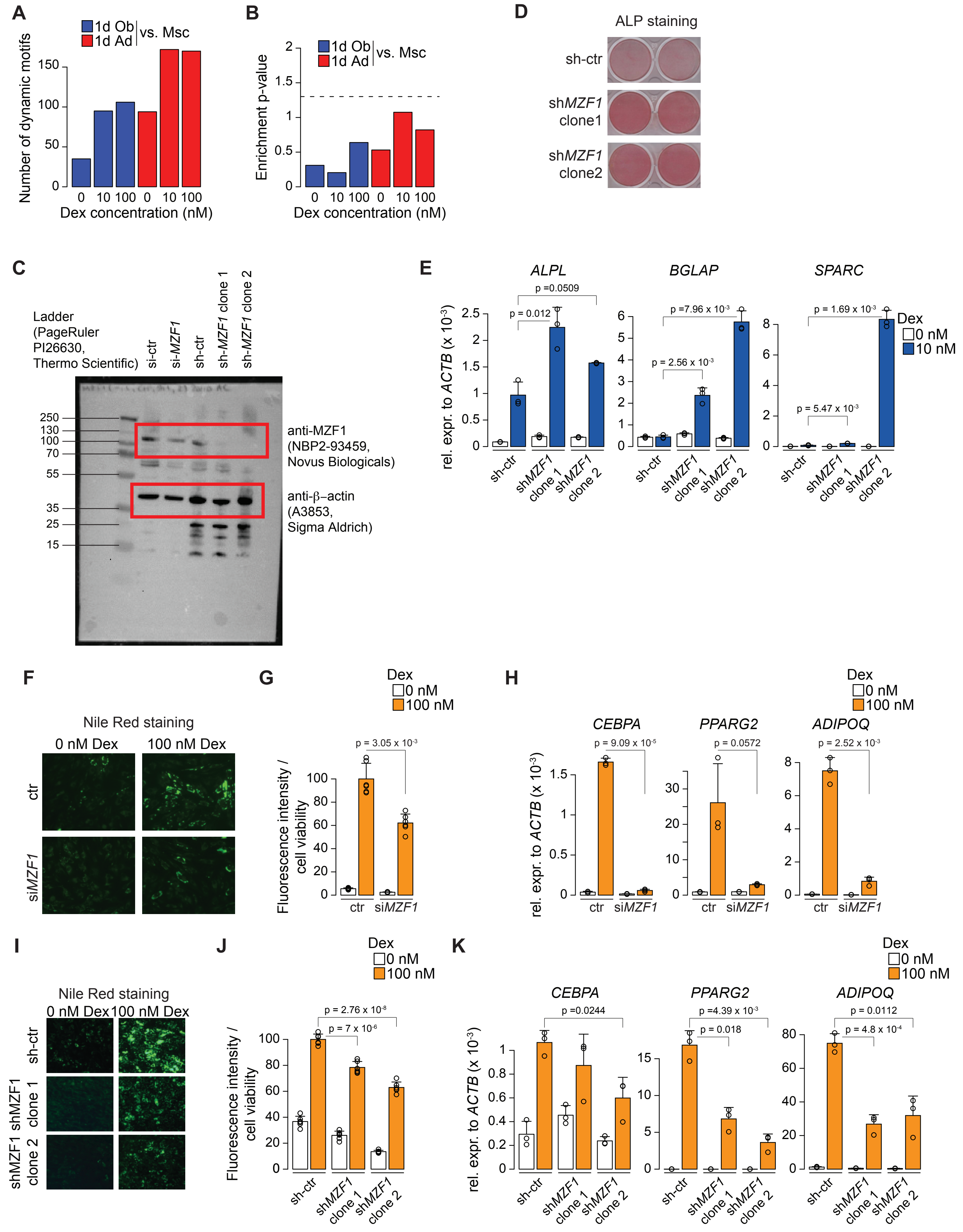

**Supplementary Figure 4 – related to Figure 4.** (**A**) Bar plot quantifying the number of motifs with differential activity (p-value < 0.05) based on H3K27ac-ChIP-seq on day 1 of osteoblast or adipocyte differentiation in hBM-MSC-TERT4 cells with indicated concentrations of Dex. (**B**) Bar plot showing the enrichment of dynamic motifs from Fig. S4A for TFs with differential expression (DESeq2 p-adj < 0.01). Enrichment based on hypergeometric test with dashed line indicating a threshold of 0.05. (**C**) Uncropped Western blot analysis shown in Fig. 4E. Membrane was used for parallel detection of MZF1 (top) and b-actin (right). Red boxes indicate the cropped region shown in Fig. 4E. (**D**) ALP staining on day 7 of osteogenic differentiation in hBM-MSC-TERT4 cells stably carrying control or shRNA constructs targeting *MZF1*. (**E**) RT-qPCR-based mRNA expression levels of *ALPL*, *BGLAP*, and *SPARC* after 7 days of osteogenic differentiation in the presence or absence of Dex in hBM-MSC-TERT4 cells stably carrying control or shRNA constructs targeting *MZF1* Bar showing mean and error bars depicting standard deviation, n = 3. Unpaired Student's t-test. (**F**) Fluorescence image of Nile Red staining in hBM-MSC-TERT4 cells after 14 days of adipocyte differentiation with or without of Dex. Cells were treated with control or *MZF1* targeting siRNA 3 days prior to induction of differentiation. (**G**) Nile Red staining intensity normalized to cell viability from pictures in Fig. S4F. Bar showing mean and error bars depicting standard deviation, n = 4. Unpaired Student's t-test. (**H**) RT-qPCR-based mRNA expression levels of *CEBPA*, *PPARG2*, and *ADIPOQ* in hBM-MSC-TERT4 cells after 7 days of adipogenic differentiation in the presence or absence of Dex. Cells were treated with control or *MZF1* targeting siRNA 3 days prior to induction of differentiation. Bar showing mean and error bars depicting standard deviation, n = 3. Unpaired Student's t-test. (**I**) Fluorescence image of Nile Red staining in hBM-MSC-TERT4 cells stably carrying control or shRNA constructs targeting *MZF1* after 14 days of adipocyte differentiation with or without Dex. (**J**) Nile Red staining intensity normalized to cell viability from pictures in Fig. S4I. Bar showing mean and error bars depicting standard deviation, n = 4. Unpaired Student's t-test. (**K**) qPCR-based mRNA expression levels of *CEBPA*, *PPARG2*, and *ADIPOQ* after 7 days of adipogenic differentiation in the presence or absence of Dex in hBM-MSC-TERT4 cells stably carrying control or shRNA constructs targeting *MZF1*. Bar showing mean and error bars depicting standard deviation, n = 3. Unpaired Student's t-test.

#### Supplementary Figure 5

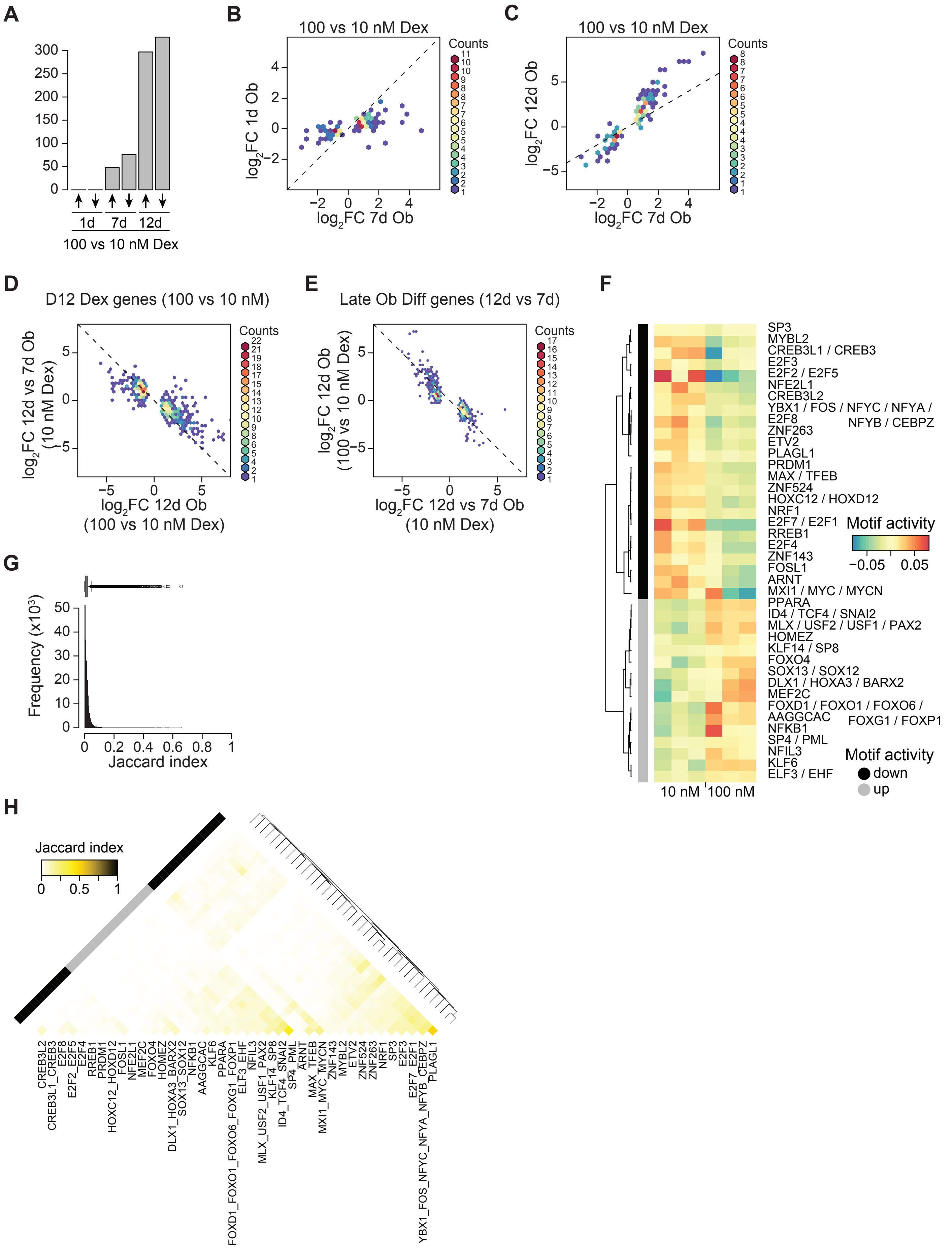

**Supplementary Figure 5 – related to Figure 5.** (**A**) Bar plot quantifying the number of differentially expressed genes (DESeq2 p-adj < 0.01) contrasting 100 versus 10 nM Dex on days 1, 7, and 12 of osteoblast differentiation. (**B**) Density scatter plot comparing changes in gene expression by 100 versus 10 nM Dex onday 7 and 1 of osteoblast differentiation. Genes selected based on differential expression by Dex on day 7 (DEseq2 padj < 0.01). Dashed line represents identity line. (**C**) Density scatter plot comparing changes in gene expression by 100 versus 10 nM Dex on days 12 and 7 of osteoblast differentiation. Genes selected based on differential expression by Dex on day 7 (DEseq2 padj < 0.01). Dashed line represents identity line. (**D**) Density scatter plot comparing changes in gene expression by 100 versus 10 nM Dex on day 12 with the transition from day 7 to day 12 of osteoblast differentiation. Genes selected based on differential expression by Dex on day 12 (DEseq2 padj < 0.01). Dashed line represents inverse identity line. (**E**) Density scatter plot comparing changes in gene expression by transition from day 7 to day 12 with 100 versus 10 nM Dex on day 12 of osteoblast differentiation. Genes selected based on differential expression between day 12 and 7 (DEseq2 padj < 0.01). Dashed line represents inverse identity line. (**F**) Heat map visualizing the ISMARA-based motif activity on day 12 of osteoblast differentiation with 100 or 10 nM Dex. Motifs selected based on differential activity (ISMARA z-score > 2) and hierarchically clustered based on activity profiles. (**G**) Histogram and box plot visualizing the distribution of pairwise Jaccard indices as a measure of overlap between predicted target gene sets of ISAMRA motifs. (**H**) Heat map visualizing pairwise Jaccard indices as an overlap between predicted target gene sets of motifs with differential activity (ISMARA z-score > 2) grouping as in Fig. S5F.

### Supplementary Figure 6

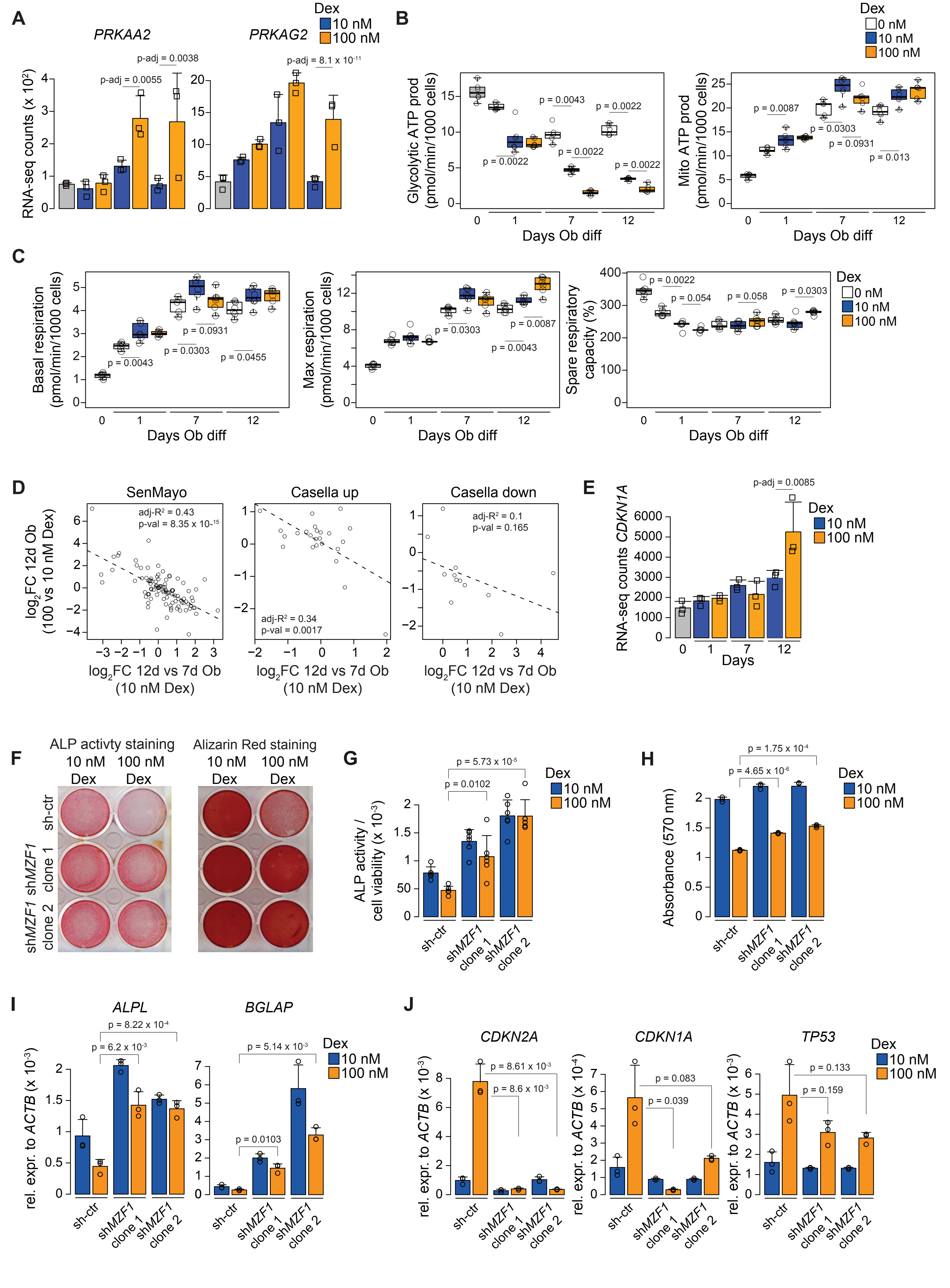

**Supplementary Figure 6 – related to Figure 6.** (**A**) RNA-seq based counts of *PRKAA2 and PRKAG2* (encoding regulatory subunits of AMPK) in hBMSC-TERT4-cells prior to and after 1, 7, or 12 days of osteogenic differentiation with 10 or 100 nM of Dex. DESeq2 adjusted p-value. (**B**) Box plot quantifying Seahorse XF analysis-based quantification of glycolytic ATP production (calculated from ECAR, left panel) and mitochondrial ATP production (calculated from OCR, right panel) with the indicated concentrations of Dex and at time points of osteogenic differentiation. Two-tailed Wilcoxon–Mann–Whitney testing Dex effect against 10 nM at a given time point. (**C**) Box plot quantifying Seahorse XF analysis-based basal respiration (left panel), maximum respiration (middle panel), and spare respiratory capacity (right panel) with the indicated concentrations of Dex and at time points of osteogenic differentiation. Two-tailed Wilcoxon–Mann–Whitney testing Dex effect against 10 nM at a given time point. (**D**) Scatter plots comparing changes in gene expression by 100 versus 10 nM Dex on day 12 with the transition from day 7 to day 12 of osteoblast differentiation. Genes selected based on senescence gene signatures SenMayo (left panel), Casella upregulated (middle panel), and Casella downregulated (right panel). Line based on linear regression with residuals. (**E**) RNA-seq based counts of *CDKN1A* (encoding p21) in hBMSC-TERT4-cells prior to and after 1, 7, or 12 days of osteogenic differentiation with 10 or 100 nM of Dex. DESeq2 adjusted p-value. (**F**) ALP (day 7) and Alizarin Red (day 14) staining upon osteogenic differentiation with 10 or 100 nM of Dex in hBM-MSC-TERT4 cells stably carrying control or shRNA constructs targeting *MZF1*. (**G**) Bar plot quantifying ALP activity normalized to cell viability in cells in Fig. S6F on day 7 of osteoblast differentiation. Bar showing mean and error bars depicting standard deviation, n = 6. Unpaired Student's t-test. (**H**) Bar plot quantifying Alizarin Red staining intensity in cells in Fig S6F on day 14 of differentiation. Bar showing mean and error bars depicting standard deviation, n = 3. Unpaired Student's t-test. (**I**) RT-qPCR-based mRNA expression levels of *ALPL* and *BGLAP* after 7 days of osteogenic differentiation with 10 or 100 nM of Dex in hBM-MSC-TERT4 cells stably carrying control or shRNA constructs targeting *MZF1*. Bar showing mean and error bars depicting standard deviation, n = 3. Unpaired Student's t-test. (**J**) RT-qPCR-based mRNA expression levels of *CDKN2A* (encoding p16), *CDKN1A* (encoding p21), and *TP53* (encoding p53) after 7 days of osteogenic differentiation with 10 or 100 nM of Dex in hBM-MSC-TERT4 cells stably carrying control or shRNA constructs targeting *MZF1*. Bar showing mean and error bars depicting standard deviation, n = 3. Unpaired Student's t-test.
